# A combinatorial lipid code shapes the electrostatic landscape of plant endomembranes

**DOI:** 10.1101/278135

**Authors:** Matthieu Pierre Platre, Mehdi Doumane, Vincent Bayle, Mathilde Laetitia Audrey Simon, Lilly Maneta-Peyret, Laetitia Fouillen, Thomas Stanislas, Laia Armengot, Přemysl Pejchar, Marie-Cécile Caillaud, Martin Potocký, Alenka Čopič, Patrick Moreau, Yvon Jaillais

## Abstract

Membrane surface charge is critical for the transient, yet specific recruitment of proteins with polybasic regions to certain organelles. In all eukaryotes, the plasma membrane (PM) is the most electronegative compartment of the cell, which specifies its identity. As such, membrane electrostatics is a central parameter in signaling, intracellular trafficking and polarity. Here, we explore which are the lipids that control membrane electrostatics using plants as a model. We show that phosphatidic acidic (PA), phosphatidylserine (PS) and phosphatidylinositol-4-phosphate (PI4P) are separately required to generate the electrostatic signature of the plant PM. In addition, we reveal the existence of an electrostatic territory that is organized as a gradient along the endocytic pathway and is controlled by PS/PI4P combination. Altogether, we propose that combinatorial lipid composition of the cytosolic leaflet of cellular organelles not only defines the plant electrostatic territory but also distinguishes different compartments within this territory by specifying their varying surface charges.

## Introduction

An evolutionarily conserved feature of cellular organelles is the distinct phospholipid composition of their membranes, which is essential to specify their identity and function. Within the endomembrane system of eukaryotic cells, the existence of two major lipid territories has been postulated, one characterized by membranes with lipid packing defects, and the other defined by membrane surface charges (Bigay and Antonny, 2012). These two lipid territories correspond roughly to two dynamic membrane-recycling systems; one centered on the endoplasmic reticulum (ER) and that includes membranes from the ER, the nuclear envelope and the *cis-*Golgi, and the other centered on the plasma membrane (PM) and that comprises the *trans*-Golgi, the *trans*-Golgi Network (TGN), the PM and endosomes (Jackson et al., 2016). In the later, referred to as the electrostatic territory, negative charges carried by anionic phospholipids recruit proteins with polybasic regions and as such participate in the localization of a large number of cellular factors along the endocytic pathway (Jackson et al., 2016). Anionic phospholipids are minor lipids in membranes and include phosphatidylinositol phosphate (also known as phosphoinositides), as well as phosphatidic acid (PA) and phosphatidylserine (PS). In mammalian cells, PS is enriched in PM-derived organelles and was proposed to act as a landmark of the electrostatic territory (Bigay and Antonny, 2012; Jackson et al., 2016; Yeung et al., 2008). However, this model was only tested *in vitro* in cultured human cells, notably in macrophages (Yeung et al., 2008; Yeung et al., 2009), and was not yet challenged in loss-of-function experiments with genetic and/or pharmacological depletion of cellular PS. In addition, it is currently unknown whether this model can be extended to other eukaryotic systems, such as fungi or plants.

A second characteristic of the electrostatic territory lays in the finding that it is not uniformly organized across all PM-derived organelles (Platre and Jaillais, 2017; Yeung et al., 2006). Rather, the inner leaflet of the PM is the most electronegative cytosolic-facing membrane across eukaryotes, including yeasts, animals and plants (Platre and Jaillais, 2017). This PM electrostatic signature is critical for cell signaling as it enables to specifically recruit proteins to the PM., such as e.g., small GTPases, kinases, or kinase regulators (Barbosa et al., 2016; Heo et al., 2006; Moravcevic et al., 2010; Noack and Jaillais, 2017; Simon et al., 2016; Yeung et al., 2006). In most eukaryotic cells, phosphoinositides species act redundantly to maintain the PM electrostatic signature, as the loss of one phosphoinositide has little or no impact on the overall charge of the membrane. For example in animal cells, acute depletion of PI(4,5)P_2_ has no effect on the PM electrostatic field, since charges from PI4P and PI(3,4,5)P_3_ are sufficient to maintain PM electrostatics (Hammond et al., 2012; Heo et al., 2006). Similarly in yeast, concomitant inhibition of both PI4P and PI(4,5)P_2_ synthesis does not impact PM electrostatics significantly (Moravcevic et al., 2010). By contrast, we recently established that PI4P is the main phosphoinositide required to power the high electrostatic field of the plant PM, while PI(4,5)P_2_ is dispensable for PM surface charges (Simon et al., 2016). The unique role of PI4P as the main phosphoinositide involved in PM electrostatics in plants raises the question of whether other anionic phospholipid species may contribute or not to the particular PM electrostatic signature.

Here, we addressed whether an electrostatic territory exists beyond the PM in plant cells and asked whether this electrostatic landscape shares features with that of animal cells. To do so, we studied the contribution of other anionic phospholipids to membrane surface charges and how they may cooperate with PI4P to regulate the establishment of an electrostatic territory in plants. We therefore focused on PS, which is thought to be important for the electrostatic landscape of animal intracellular compartments (Bigay and Antonny, 2012). PA is not normally present at the PM in animal cells (Bohdanowicz et al., 2013), but has been visualized at the PM in plant tip growing cells (Noack and Jaillais, 2017; Potocky et al., 2014). In addition, PA is present at relatively high level in plant cells (e.g. 5% of total phospholipid in rye seedling (Lynch and Steponkus, 1987)). We therefore also studied the potential role of PA in setting up PM electrostatics. To address whether PA and PS could contribute to the establishment of an electrostatic territory, we first analyzed their subcellular localization using genetically encoded biosensors. We further used these sensors to validate pharmacological and genetic approaches designed to conditionally perturb the production of these lipids. We demonstrate that PA and PS act in concert with PI4P to generate the distinctively high PM electrostatic field. In addition, we reveal the existence of an electrostatic gradient along the endocytic pathway, being the highest at the PM, intermediate in early endosomes/*trans*-Golgi Network (EE/TGN) and lowest in late endosomes (LE). We further show that PS, in combination with PI4P, organizes this intracellular electrostatic gradient. Overall, we demonstrate the existence of an electrostatic territory that corresponds to PM-derived organelles in plants and resembles that of animal cells. In addition, we show that within this territory, each compartment has a distinct electrostatic signature that is set-up by a combinatorial code of various anionic phospholipids. We propose that this “electrostatic code” may represent a fundamental patterning principle of the endomembrane system and acts as a key determinant of protein subcellular targeting.

## Results

### PA accumulates at the PM cytosolic leaflet in *Arabidopsis* root epidermis

PA is an anionic phospholipid, which accumulates in the sub apical region of the PM cytosolic leaflet in tobacco pollen tubes (Potocky et al., 2014). To analyze whether PA could also localize at the PM in *Arabidopsis* sporophytic tissues, and thereby may contribute to PM electrostatics, we raised transgenic *Arabidopsis* lines stably expressing mCITRINE-tagged variants of the recently developed “*PA biosensor with superior sensitivity*” (mCITRINE-1xPASS and mCITRINE-2xPASS) (Lu et al., 2016; Zhang et al., 2014) under the control of the mild ubiquitous promoter of the *UBIQUITIN10* (*UBQ10*) gene. This PA probe is based on the PA-binding motif of the yeast Spo20p protein, with an extra nuclear export signal (NES) to exclude the fusion protein from the nucleus and increase the accessibility of the probe to the cytosol (Lu et al., 2016; Zhang et al., 2014). Both mCITRINE-1xPASS and mCITRINE-2xPASS sensors were targeted to the PM in *Arabidopsis*, including root and shoot tissues (Figure 1A, Figure S1A and S1B). We noticed that these PA probes localized early at the cell plate (Video S1) and colocalized with the endocytic dye FM4-64 (Figure S1C), one of the earlier marker incorporated into the membrane of this compartment (Dettmer et al., 2006). Furthermore, mCITRINE-2xPASS localized on the flank region of growing root hairs (Video S2), in a pattern that closely resembled the localization of PA sensor in growing tobacco pollen tubes (Potocky et al., 2014). While both PA sensors localized at the PM, mCITRINE-1xPASS was also cytosolic, while mCITRINE-2xPASS accumulated in the nucleus (Figure 1A). This suggests that in mCITRINE-2xPASS, the NES is not as efficient as in the mCITRINE-1xPASS probe. Consistently, the mCITRINE-1xPASS probe for which the NES is mutated (1xPASS^NESmut^) localized at the PM, the cytosol and in the nucleus (Figure 1A). It is unclear what the significance of the nuclear localization of the probe is. Indeed, it might reflect uncontrolled diffusion from the cytosol into the nucleus or trapping of the probe in the nucleus by nuclear PA. Of note, for all three transgenic lines (mCITRINE-1xPASS, mCITRINE-1xPASS^NESmut^, mCITRINE-2xPASS), we observed some variability on the intensity of PM labeling between different roots. Although the cause of this variability is currently unknown, it might arise from different stress status of individual roots or cells since PA metabolism is well known to be under tight environmental control (Testerink and Munnik, 2011). Nonetheless, the three aforementioned probes are targeted to the PM of root meristematic cells (Figure 1A), suggesting local enrichment of PA in this membrane even in normal growing conditions (i.e. non-stressed).

**Figure 1.**
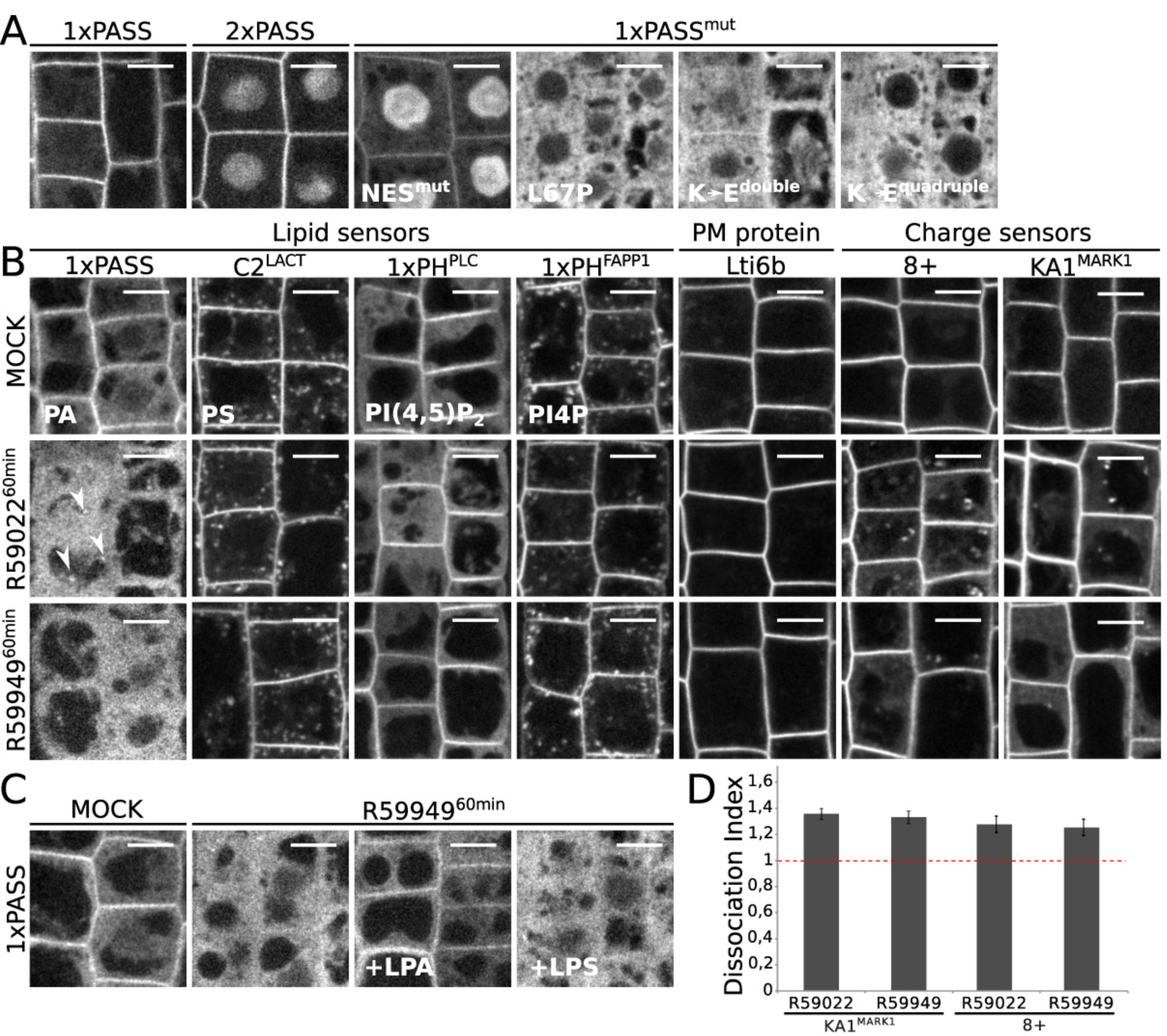
DAG Kinase-dependent accumulation of phosphatidic acid at the PM is required to maintain the electrostatic field of the PM cytosolic leaflet. **A**, Confocal images of *Arabidopsis* root epidermis expressing from left to right, mCITRINE-1xPASS, mCITRINE-2xPASS, mCITRINE-1xPASS^NESmut^, mCITRINE-1xPASS^L^67^P^, mCITRINE-1xPASS^K^66^E-K^68^E^ (K→E double), and mCITRINE-1xPASS^K^66^E-K^68^E-^ K71E-K73E (K→E quadruple). **B**, Confocal images of plants expressing from left to right, PA, PS, PI(4,5)P_2_ and PI4P sensors (mCITRINE-1xPASS, mCITRINE-C2^LACT^, mCITRINE-1xPH^PLC^, mCITRINE-1xPH^FAPP1^), plasma membrane-associated protein (EGFP-Lti6b) and charge sensors (mCITRINE^8K-Farn^ (8+), mCITRINE-KA1^MARK1^), in mock conditions (top), plants treated with 12.5µM R59022 (middle) or 12.5µM R59949 (bottom) for 60 min. Arrows highlight the presence of spots. **C**, Confocal images of *Arabidopsis* root epidermis expressing mCITRINE-1xPASS upon concomitant lysoPA (LPA) or lysoPS (LPS) and R59949(12.5 µM) treatment for 60min. From left to right, mock, R59949alone, R59949+ LPA, and R59949+ LPS. **D,** Quantification of the mCITRINE-KA1^MARK1^ and mCITRINE^8K-Farn^ dissociation index (mean ±s.e.m), upon R59022 and R59949 treatment (n=150 cells 12.5µM, 60min). Scale bars, 5 µm.

The PA-binding motif of Spo20p was extensively validated as a sensor *in vivo* in animal cells (Bohdanowicz et al., 2013; Zhang et al., 2014), as well as in pollen tube (Potocky et al., 2014). However, *in vitro*, PA binding was also shown to be dependent on the local lipid environment of the probe (i.e. local surface charges) (Horchani et al., 2014; Kassas et al., 2017). In order to validate the PA sensor specificity *in planta*, we first expressed mCITRINE-1xPASS mutant versions (L67P single mutant, K66E-K68E double mutants and K66E-K68E-K71E-K73E quadruple mutants), which were previously shown to impair PA binding (Potocky et al., 2014). mCITRINE-1xPASS^K66E-K^68^E^ (1xPASS^K>E^ ^double^) retained a faint PM labeling, while mCITRINE-1xPASS^L67P^ and mCITRINE-1xPASS^K66E-K68^E-K^71E-K^73^E^ (1xPASS^K>E^ ^quadruple^) were fully soluble, suggesting that lipid binding is required for the PM localization of the 1xPASS probe (Figure 1A). Diacylglycerol kinases (DGK) are the major PA producing enzymes at the PM of animal cells with constitutively elevated PA level (Bohdanowicz et al., 2013). We therefore analyzed the effect of R59949 and R59022, two inhibitors of DGK activity, on the localization of PA reporters. Both inhibitors induced the release of mCITRINE-1xPASS and mCITRINE-2xPASS PA probes from the PM into the cytosol and nucleus, respectively (Figure 1B and S1D). These results suggest that DGKs are required to maintain PA production at the plant PM. To confirm that the dissociation of mCITRINE-1xPASS was caused by inhibition of PA production in R59949 treated seedling, we performed add-back experiments by supplementing the root with exogenous lysophosphatidic acid (LPA) or lysophosphatidylserine (LPS) as control. We used lysophospholipids since they have identical head groups as PA/PS but are more soluble than phospholipids and as such are more likely to reach the cytosolic leaflet of cellular membranes (Moser von Filseck et al., 2015a). We found that upon inhibition of endogenous PA production by R59949, mCITRINE-1xPASS was maintained at the PM in presence of an exogenous supply of LPA but not in the presence of LPS (Figure 1C). Moreover, in the presence of either R59949 or R59022, reporters for PI4P, PI(4,5)P_2_ and PS anionic phospholipids (mCITRINE-1xPH^FAPP1^, mCITRINE-1xPH^PLC^ and mCITRINE-C2^Lact^ respectively (Simon et al., 2014; Simon et al., 2016)), were still localized at the PM (Figure 1B). Altogether, these results indicated that the PM localization of the mCITRINE-1xPASS and mCITRINE-2xPASS probes are largely driven by DGK-synthesized PA, rather than by a general requirement of these probes for anionic phospholipids. In addition, both R59949 and R59022 treatments had no impact on the localization of EGFP-Lti6b (Figure 1B), a control protein with two transmembrane segments and very short cytosolic tails, whose localization is not regulated by anionic lipids (Cutler et al., 2000). Altogether, these results validate the specificity of our PA probes and suggest that PA accumulates in the cytosolic leaflet of the plant PM. It is therefore possible that this anionic lipid participates in the control of PM electrostatics.

### PA contributes to PM cytosolic leaflet surface charges

We next asked whether PA could participate in the electrostatic property of the PM. We took advantage of DGK inhibitors to reduce the level of PA at the PM and analyze the impact of this pharmacological inhibition on the localization of membrane surface charge reporters. We used two types of membrane charge reporters that we previously validated *in planta* (Simon et al., 2016). The first probe, mCITRINE^8K-Farn^ corresponds to two mCITRINE fluorescent proteins fused in tandem, which localize in electrostatic membranes thanks to the combinatorial effects of a polycationic region (with 8 net positive charges, +8) and an adjacent farnesyl lipid anchor, which provides hydrophobic anchoring (Haupt and Minc, 2017; Platre and Jaillais, 2017; Simon et al., 2016; Yeung et al., 2008; Yeung et al., 2006). The second probe corresponds to the KA1 domain of the human protein MARK1, which is a folded unit known to interact non-stereospecifically with all anionic phospholipids (Hammond et al., 2012; Moravcevic et al., 2010; Platre and Jaillais, 2017; Simon et al., 2016). We found that in PA depleted condition, charge sensors (mCITRINE^8K-Farn^ and mCITRINE-KA1^MARK1^ probes) were released into the cytosol and endosomes (Figure 1B and D). Endosome labelling was more prominent with R59022 than R59949 treatment and correlated with the concomitant accumulation of mCITRINE-1xPASS in similar compartments (see arrows, Figure 1B). Together, our results suggest that PA contributes to the electrostatic properties of the plasmalemma cytosolic leaflet.

### PS accumulates on the cytosolic leaflet of PM and PM-derived organelles

To evaluate the potential function of PS in membrane electrostatics, we studied its sub-cellular distribution using genetically encoded biosensors that report the localization of PS in inner membrane leaflets. We used the stereospecific PS-binding C2 domain of bovine Lacthaderin (C2^LACT^) and the pleckstrin homology (PH) domain of human EVECTIN2 (PH^EVCT2^). These probes were extensively validated as calcium-independent PS reporters (Chung et al., 2015; Haupt and Minc, 2017; Moravcevic et al., 2010; Moser von Filseck et al., 2015a; Simon et al., 2016; Uchida et al., 2011; Yeung et al., 2008; Yeung et al., 2009) (Figure S2A and S2B). We raised transgenic *Arabidopsis* plants that stably express fluorescent fusions with either C2^LACT^, or 2xPH^EVCT2^ under the control of the *UBQ10* promoter. As we previously reported for mCITRINE-C2^LACT^ in root epidermis (Simon et al., 2016), we found that the C2^LACT^ domain was localized at the PM and in multiple intracellular compartments in all cell types analyzed, including both shoot and root tissues (Figure 2A and Figure S2C-J). We noticed that in tip growing cells such as root hairs and pollen tubes, C2^LACT^, was localized to the shank region of the plasma membrane and intracellular compartments and accumulated in the inverted cone region at their very tip (Video S3 and S4, respectively), a region known for active endocytic and exocytic activities (Noack and Jaillais, 2017). In addition, the mCITRINE-2xPH^EVCT2^ reporter showed a similar localization pattern as mCITRINE-C2^LACT^ and, consistently, tdTOMATO-2xPH^EVCT2^ extensively colocalized with mCITRINE-C2^LACT^ (Figure S2C). However, similarly to animal cells (Chung et al., 2015; Uchida et al., 2011; Yeung et al., 2008), we noticed that mCITRINE-C2^LACT^ PM localization was more pronounced than that of mCITRINE-2xPH^EVCT2^/tdTOMATO-2xPH^EVCT2^.

**Figure 2.**
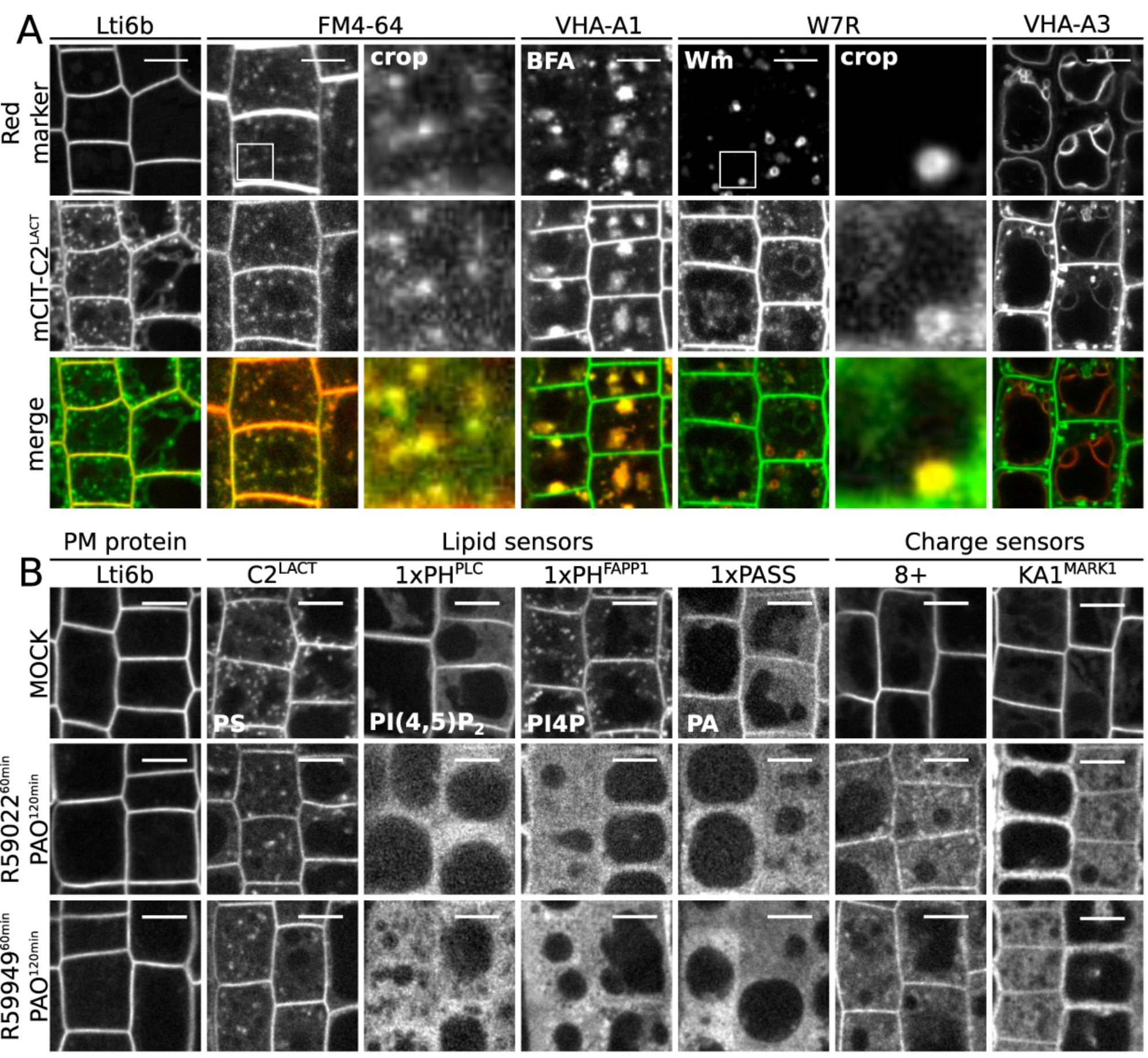
Phosphatidylserine accumulates at the PM and along the endocytic pathway and is sufficient to maintain negative charges at the PM cytosolic leaflet. **A)** Confocal images of *Arabidopsis* root epidermis co-expressing a red fluorescence marker (top), mCITRINE-C2^LACT^ (middle), and corresponding merge (bottom). Top images correspond to (from left to right): Lti6b-2xmCHERRY (PM marker), FM4-64 (endocytic tracer, 1µM, 60 min), VHA-A1-mRFP1 (EE/TGN marker) in the presence of brefeldinA (BFA, 25µM, 60min), W7R (LE marker) treated with 30µM wortmannin (Wm, 30µM, 90min), VHA-A3-mRFP1 (tonoplast marker). **B,** Confocal images of plants expressing from left to right, EGFP-Lti6b, mCITRINE-C2^LACT^ (PS), mCITRINE-1xPH^PLC^ (PI(4,5)P_2_), mCITRINE-PH^FAPP1^ (PI4P), mCITRINE-1xPASS (PA), mCITRINE^8K-Farn^ (membrane charge) and mCITRINE-KA1^MARK1^ (membrane charge), in mock conditions (top), plants pre-treated with 30µM PAO for 60 min and then concomitantly treated with 12.5µM R59022 and 30µM PAO for 60 min (middle), plants pre-treated with 30µM PAO for 60 min and then concomitantly treated with 12.5µM R59949and 30µM PAO for 60 min (bottom). Scale bars, 5 µm.

Next, we analyzed in which endomembrane structures the C2^LACT^ probe localized. We crossed the mCITRINE-C2^LACT^ reporter line with various red-fluorescent membrane markers lines or imaged it in conjunction with red-fluorescent dyes (Figure 2A). mCITRINE-C2^LACT^ extensively colocalized with the plasmalemmal marker Lti6b-2xmCHERRY (Elsayad et al., 2016), confirming that this PS sensor accumulates at the PM (Figure 2A). We also found that mCITRINE-C2^LACT^ was localized along the endocytic pathway. Indeed, mCITRINE-C2^LACT^ colocalized with the endocytic tracer FM4-64 and its localization was sensitive to both the fungal toxin brefeldinA (BFA) and wortmannin (Wm, Figure 2A), two drugs that affects the morphology of early and late endocytic compartments, respectively (Bayle et al., 2017; Dettmer et al., 2006; Geldner et al., 2009; Jaillais et al., 2008; Jaillais et al., 2006). Finally, we observed in few meristematic cells (14.3% s.e.m. ±2.73, n=458 cells) that mCITRINE-C2^LACT^ colocalized with the tonoplast marker VHA-A3-mRFP1 (Figure 2A) (Dettmer et al., 2006). We also found that mCITRINE-C2^LACT^ localized early on forming cell plate during cytokinesis, together with FM4-64 and PI4P (Figure S2K and Video S5). Therefore, PS accumulation at the cell plate together with PA and PI4P correlates with the acquisition of the cell plate electrostatic identity (Simon et al., 2016). Together, our results suggest that PS accumulates at the PM and cell plate, as well as in PM-derived organelles.

### PS is sufficient to maintain negative surface charges on the PM cytosolic leaflet

Next, we addressed whether PS contributes to PM electrostatics. Because there is no chemical compound known to directly inhibit PS production, we tested whether PS could be involved in PM electrostatics by depleting all other anionic phospholipids from this membrane through chemical inhibition. We previously validated the use of PAO, a PI4-Kinase inhibitor, to interfere with PM phosphoinositides production (Simon et al., 2016). We showed that short-term treatment (15-30 min) significantly depletes PI4P but not PI(4,5)P_2_ pools, while longer treatment (>60 min) affects the synthesis of both lipids (Simon et al., 2016). In order to concomitantly deplete the plant PM from PA, PI4P and PI(4,5)P_2_, leaving PS as the sole anionic lipid in this membrane, we used a combination of R59949 or R59022 (60 min, as described in Figure 1) and prolonged PAO treatment (120 min). This treatment efficiently displaced PI4P, PI(4,5)P_2_ and PA sensors from the PM to the cytosol, while the PM localization of EGFP-Lti6b and mCITRINE-C2^LACT^ were largely unaffected by this treatment (Figure 2B). As expected, a proportion of mCITRINE^8K-Farn^ and mCITRINE-KA1^MARK1^ charge reporters were found in the cytosol in this condition (Figure 2B). However, surprisingly, both charge reporters retained a degree of PM localization that can be attributed to PS, the only remaining anionic lipid in this membrane. Given the physiological importance of PA, PI4P and PI(4,5)P_2_, this concomitant treatment is expected to have pleiotropic detrimental effects on plant cell biology, notably inhibiting various intracellular trafficking pathways such as endocytosis and exocytosis as well as signaling pathways. Nonetheless, in this condition, PS appears to be sufficient to maintain a certain degree of surface charges at the PM.

### *phosphatidylserine synthase1* mutants do not produce any PS but are viable

In order to analyze the impact of PS depletion on membrane surface charges, we characterized mutants in the *PHOSPHATIDYLSERINE SYNTHASE1* (*PSS1*) gene (Yamaoka et al., 2011). We isolated three *pss1* alleles that we named *pss1-3*; *pss1-4* and *pss1-5* (Figure 3A). These three alleles expressed no detectable full length *PSS1* cDNA (Figure S3C), and segregated as single recessive mutants without any distorted segregation (Figure S3A). All three alleles showed the same sporophytic phenotype, the *pss1* mutants being severely dwarf both at the shoot and root level (Figure 3B, S3F-I). In addition, these mutants were sterile and had to be propagated as heterozygous. Next, we introduced a wild type copy of the *PSS1* gene in the *pss1-3* allele, which fully complemented the *pss1-3* shoot phenotypes (Figure 3B, S3D and F). High performance thin layer chromatography (HPTLC) and LC-MS/MS lipidomic analyses showed that *pss1-3* and *pss1-4* sporophytes do not produce any PS (Figure 3C-D and table S1). Importantly, these analyses suggested that both alleles had only minor changes in their overall phospholipid content (Figure 3C, S3B and table S1). To confirm these biochemical analyses, we introgressed mCITRINE-C2^LACT^ and mCITRINE-2xPH^EVCT2^ into the *pss1-3* mutant. By contrast to the wild type situation, we could detect only a faint signal for mCITRINE-C2^LACT^ in *pss1-3*, suggesting that in the absence of PS, mCITRINE-C2^LACT^ is unstable in plant cells (Figure 3E). Consistently, exogenously treating mCITRINE-C2^LACT^/*pss1-3*^*-/-*^ seedlings for one hour with LPS, but not LPA, fully complemented mCITRINE-C2^LACT^ fluorescence signal intensity and localization at the PM and intracellular compartments (Figure 3E). In addition, the mCITRINE-2xPH^EVCT2^ probe was fully soluble in *pss1-3*, as expected for a PS-depleted mutant and this localization was rescued by one hour LPS add back experiments but not by exogenous treatment with LPA (Figure 3E). Furthermore, both root and shoot phenotypes were partially rescued by exogenous treatment with LPS (Figure S3E-I). Together, our biochemical, cell biological and phenotypical analyses suggest that *pss1-3* and *pss1-4* mutants do not produce any PS, which seems dispensable for gametogenesis and embryonic development but is absolutely required for normal post-embryonic plant development and sporophyte fertility. In addition, this PS-depleted mutant further validates the specificity of our PS-binding probes C2^LACT^ and 2xPH^EVCT2^.

**Figure 3.**
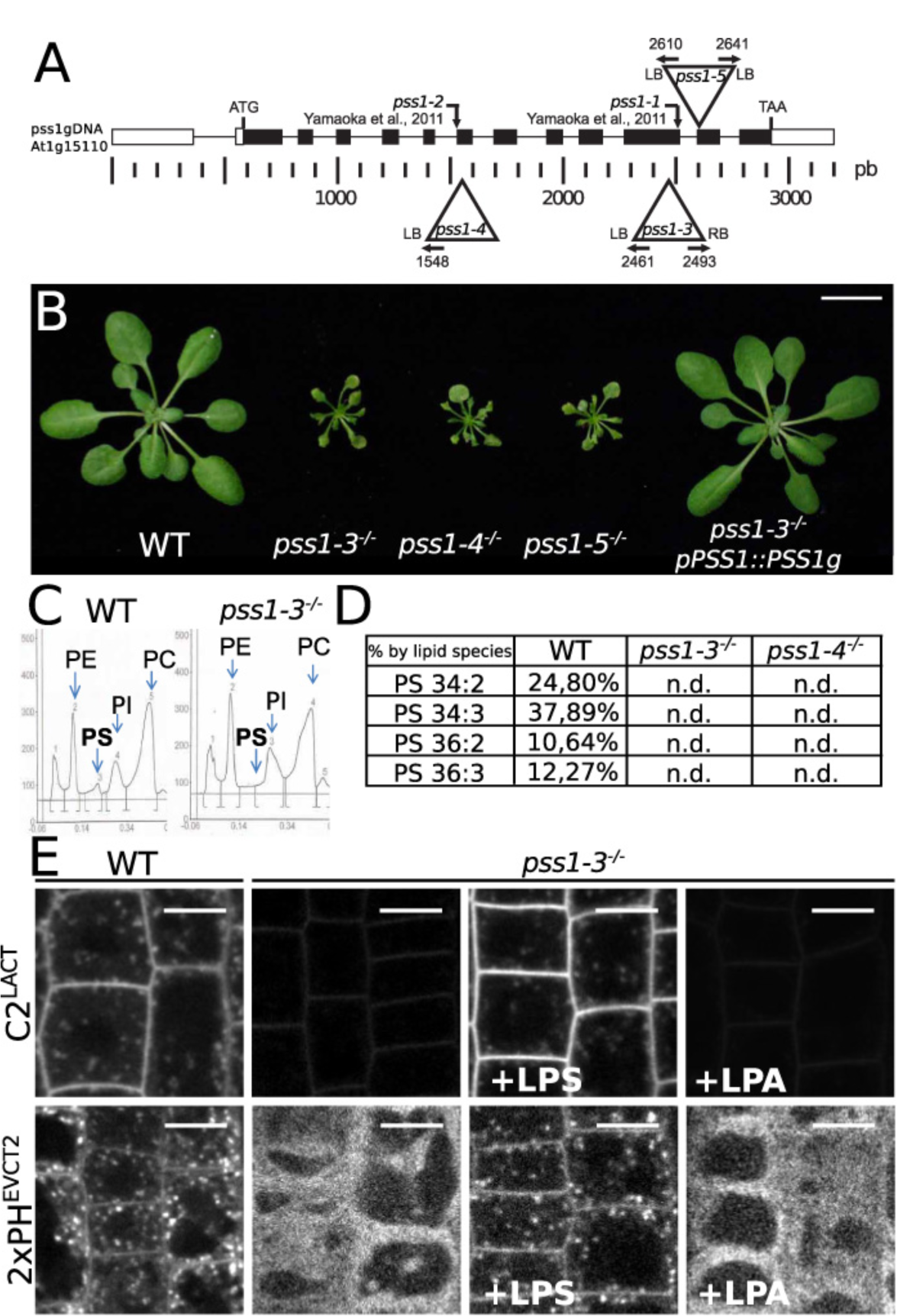
PSS1 is required for PS biosynthesis and plant growth. **A,** Schematic representation of T-DNA insertions in *PSS1*. LB, left border; RB, right border; numbers indicate the position of border/*PSS1* junctions. **B,** Rosette phenotype of *pss1* mutants compared to the wild type. From left to right, wild type (WT, Col0), *pss1-3*^*-/-*^, *pss1-4*^*-/-*^, *pss1-5*^*-/-*^ and *pss1-3*^*-/-*^ expressing *pPSS1::PSS1g*. Scale bar, 2 cm. **C,** High performance thin layer chromatography (HPTLC) assay showing a representative quantification of the phospholipids phosphatidylcholine (PC), phosphatidylethanolamine (PE), phosphatidylinositol (PI) and phosphatidylserine (PS) in WT and *pss1-3*^*-/-*^ seedlings. **D**, Table showing the percentage of the four major PS species in WT, *pss1-3*^*-/-*^ and *pss1-4*^*--/*^ quantified by LC-MS/MS. n.d., non-detected. For an extended table of the molecular composition of PC/PE/PI/PS species, see table S1. **E**, Confocal images of *Arabidopsis* root epidermis expressing mCITRINE-C2^LACT^ (top) and mCITRINE-2xPH^EVCT2^ (bottom), from left to right in WT, *pss1-3*^*-/-*^, *pss1-3*^*-/-*^ supplemented with 54µM lysoPS (LPS) or LysoPA (LPA) for 60 min. Scale bars, 5 µm.

### PS is required for surface charges of the PM cytosolic leaflet

Next, we analyzed the localization of our membrane charge reporters in *pss1-3* mutant background. Both mCITRINE^8K-Farn^ and mCITRINE-KA1^MARK1^ retained a certain degree of PM localization in *pss1-3*, but also relocalized in the cytosol and were found in intracellular compartments (Figure 4A and C). Quantification showed that the PM dissociation of mCITRINE-KA1^MARK1^ was weaker in *pss1-3* than upon PI4P depletion (i.e. PAO treatment), and similar as upon PA depletion (i.e. R59022) (Figure 4C). We next investigated whether loss of PS could be the primary cause behind these defects in PM electrostatics. First, we found that the strict PM localization of membrane charge reporters was fully restored by short-term (one hour) add-back experiment with LPS (Figure 4A and 4C). Second, since we previously showed that PI4P and PA regulate PM electrostatics, we next asked whether loss of PS might affect PM anionic phospholipid subcellular distribution. Interestingly, the PM localization of PI(4,5)P_2_, PI4P and PA sensors were not affected in *pss1-3* (Figure 4D). In addition, by introgressing in *pss1-3* various control fluorescent markers of the PM (Figure 4B) and intracellular compartments (Figure S4), we could not detect any phenotype suggesting general defects in PM protein localization, membrane organization, and/or compartments morphogenesis.

**Figure 4.**
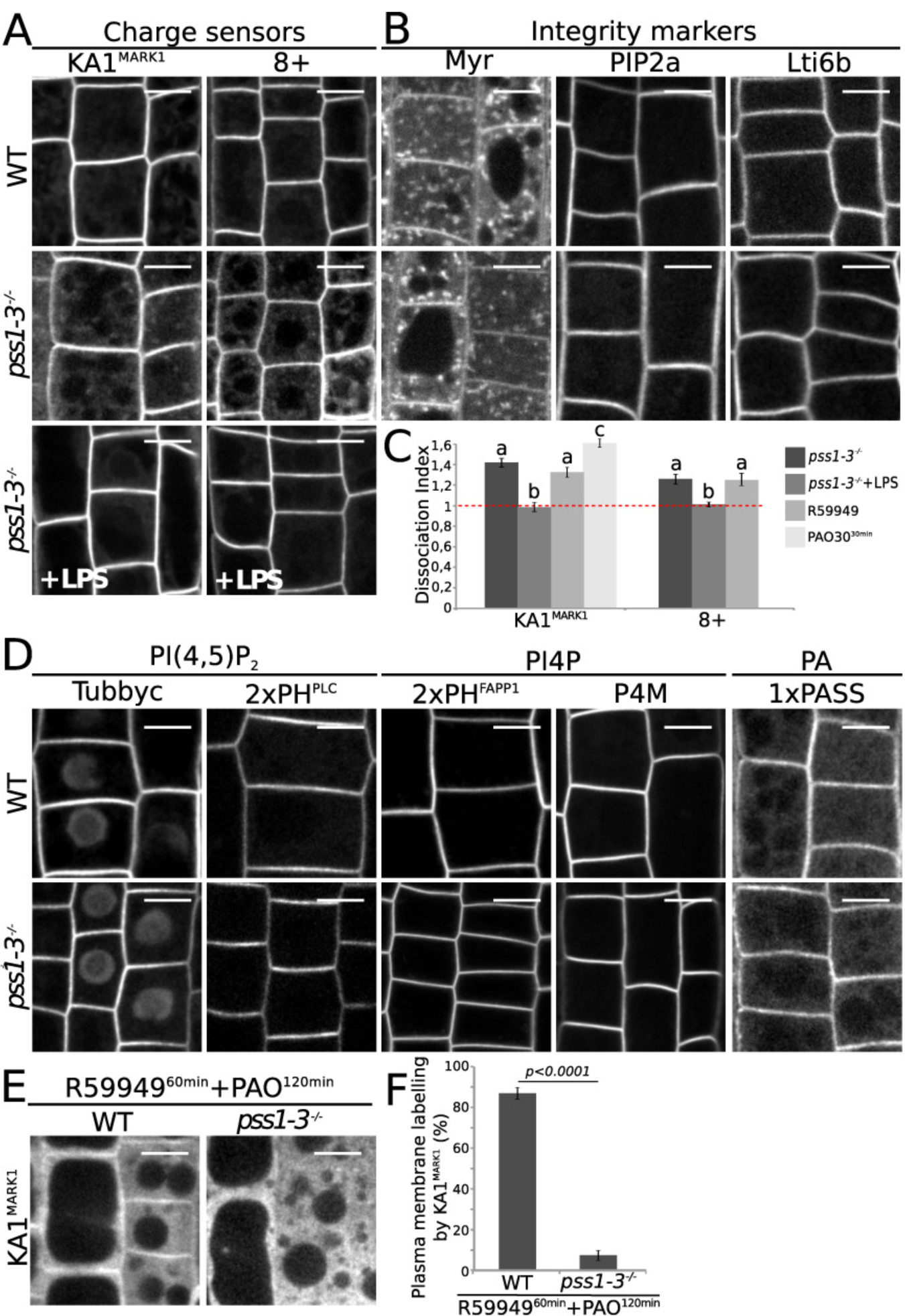
PS contributes to PM surface charges but is not required for the localization of other anionic phospholipids. **A,** Confocal images of *Arabidopsis* root epidermis expressing, mCITRINE-KA1^MARK1^ (left, KA1^MARK1^), mCITRINE^8K-Farn^ (right, 8+) in WT (top), *pss1-3*^*-/-*^ (middle), and *pss1-3*^*-/-*^ supplemented with 54µM lysoPS (LPS) for 60 min (bottom). B, Confocal images of *Arabidopsis* root epidermis expressing PM integrity markers Myr-mCITRINE (myristoylation, Myr), EGFP-PIP2a and EGFP-Lti6b in WT (top) and *pss1-3*^*-/-*^ (bottom). **C**, Quantification (mean ±s.e.m, n=150 cells) of mCITRINE-KA1^MARK1^ (left) and mCITRINE-8K^Farn^ (8+, right) dissociation index in *pss1-3*^*-/-*^, *pss1-3*^*-/-*^ supplemented with 54µM LPS for 60 min, 12.5µM R59949 for 60 min (same data set as in Figure 1D) and 30µM PAO for 30 min. Different letters indicate significant differences among means (p-value=0.05, Kruskal-Wallis bilateral test) **D**, Confocal images of *Arabidopsis* root epidermis expressing from left to right, PI(4,5)P_2_ sensors (mCITRINE-TUBBY-C (P15Y) and mCITRINE-2xPH^PLC^ (P24Y)), PI4P sensors (mCITRINE-2xPH^FAPP1^ (P21Y) and mCITRINE-P4M^SidM^) and PA sensor (mCITRINE-1xPASS) in WT (top) and *pss1-3*^*-/-*^ (bottom). **E**, Confocal images of WT (left) and *pss1-3*^*-/-*^ (right) root epidermis expressing mCITRINE-KA1^MARK1^ pre-treated with 30µM PAO for 60 min and then concomitantly treated with 12.5µM R59949 and 30µM PAO for 60 min. **F,** Quantification (mean ±s.e.m) of the percentage of cells with mCITRINE-KA1^MARK1^ at the PM in WT (left, n=887cells) and *pss1-3*^*-/-*^ (right, n=806 cells) (same treatment as in E). Statistical analysis was performed using the non-parametric Wilcoxon-Mann-Whitney test (p-value=0.05). Scale bars, 5 µm.

As described above, PS is presumably the last remaining anionic phospholipid at the PM following depletion of cellular PI4P/PI(4,5)P_2_/PA using a combination of PAO and R59949 treatment. If this assumption is correct, the vast majority of anionic phospholipids should be removed from the PM in the *pss1* mutant following this treatment, which should therefore trigger a full dissociation from the PM of our mCITRINE-KA1^MARK1^ membrane charge reporter. Concomitant PAO/R59949 treatment in *pss1-3*, indeed induced a complete loss of PM localization of mCITRINE-KA1^MARK1^, which became fully soluble in the cytosol (Figure 4E and F). This experiment demonstrates that PM localization of mCITRINE-KA1^MARK1^ in wild-type plants following concomitant PAO/R59949 treatment can be attributed to PS. Altogether, our results show that PS is not directly involved in the PM localization of other anionic lipids, but contribute to PM surface charges.

### PS localization correlates with that of electrostatic compartments

Because PS was proposed to be an important component of the electrostatic territory (Bigay and Antonny, 2012; Jackson et al., 2016), we next asked whether PS could also participate in membrane surface charges of intracellular compartments. To this end, we first mapped PS intracellular localization using quantitative colocalization analyses (see Fig S5 for a description of the method). In accordance with the BFA and Wm sensitivity we previously reported (Figure 2A), both the mCITRINE-C2^LACT^ and mCITRINE-2xPH^EVCT2^ probes localized in post-Golgi/endosomal (PG/E) compartments (Figure 5A and 5B). Interestingly, we found that both PS probes accumulated according to a concentration gradient, which is higher in early endocytic compartments (including EE/TGN and secretory vesicles (SV)), intermediate in the Golgi apparatus (Golgi) and lower in late endosomes (LE) (Figure 5B).

**Figure 5.**
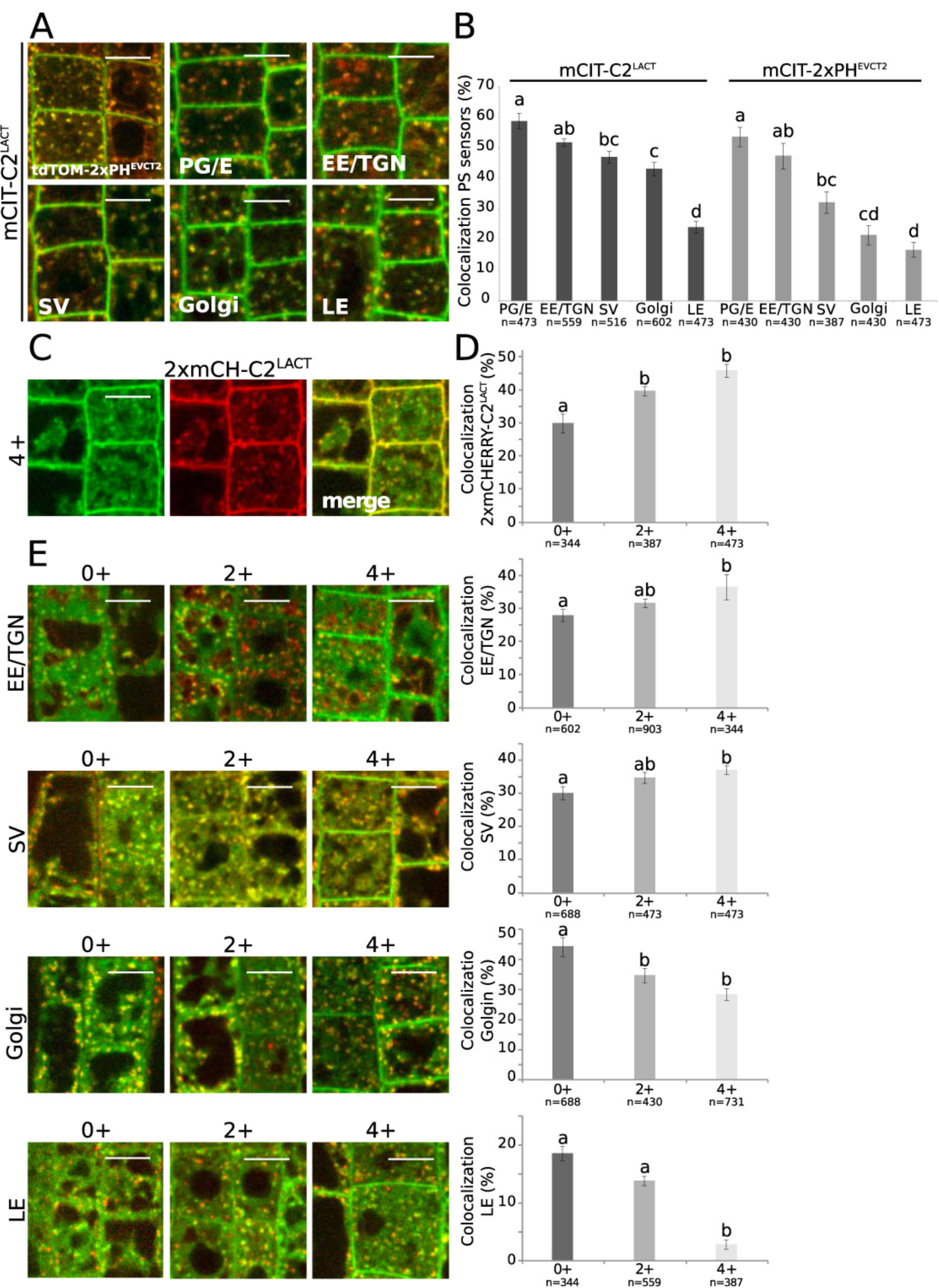
A PS gradient along the endocytic pathway correlates with a gradient of electrostatics. **A,** Merged confocal images of *Arabidopsis* root epidermis of plants co-expressing mCITRINE-C2^LACT^ with tdTOMATO-PH^EVCT2^ (top left), W25R post-Golgi endosomal/endosomes (PG/E) marker (top middle), W13R early endosomal (EE/TGN) marker (top right), W24R secretory vesicle (SV) marker (bottom left), W18R Golgi marker (bottom middle), W7R late endosomal (LE) marker (bottom right). **B,** Quantification of the percentage of compartments labelled by PS sensors (mCITRINE-C2^LACT^ and mCITRINE-2xPH^EVCT2^) that also contain compartment markers (same as above-mentioned), n=(387, 602) cells (mean ±s.e.m, percentage of colocalization). Different letters indicate significant differences among means (p value= 0.05, Kruskal-Wallis bilateral test) **C,** Confocal images of plants co-expressing mCITRINE^4K^4^Q-Farn^ (left) and 2xmCHERRY-C2^LACT^ (middle) and merge channel (right). **D**, Quantification (mean ±s.e.m) of the percentage of compartments labelled by PS sensors (2xmCHERRY-C2^LACT^) that also contain membrane charge reporters (mCITRINE^8Q-Farn^ (0+), 2xmCITRINE^2K^6^Q-Farn^ (2+) and 2xmCITRINE^4K^4^Q-Farn^ (4+)), n=(387, 602) cells. Different letters indicate significant differences among means (p value= 0.05, Kruskal-Wallis bilateral test), **E,** Merged confocal images (left) and colocalization quantification (mean ±s.e.m, right) of plants co-expressing mCITRINE^8Q-Farn^ (0+, left), mCITRINE^2K^6^Q-Farn^ (2+, middle) and mCITRINE^4K^4^Q-Farn^ (4+, right) with, from top to bottom, W13R (EE/TGN), W24R (SV), W18R (Golgi) and W7R (LE) markers, n=(344, 688) cells. Different letters indicate significant differences among means (p value= 0.15, Kruskal-Wallis bilateral test). In each graph, “n” represents the estimated number of cells sampled in each condition. Scale bars, 5 µm.

Next, we addressed which intracellular compartments were electronegative. To this end, we used charge reporters that are hydrophobically-anchored to membrane via a farnesyl moiety and that have an adjacent unstructured peptide of net varying charges (from +0 to +8) (Simon et al., 2016). A neutral version of the probe (+0, 8Q-Farn) is localized only by the intrinsic properties of the farnesyl lipid anchor, independently of membrane electrostatics. The gradual addition of positive charges by substitution of neutral glutamines into cationic lysines gradually increases the avidity of the probes for anionic membranes. As a result, a probe with intermediate charges (e.g. 4K4Q-Farn, 4+) resides in compartments that are electronegative indistinctively of whether they are highly negatively charged or not (Haupt and Minc, 2017; Platre and Jaillais, 2017; Simon et al., 2016; Yeung et al., 2008; Yeung et al., 2006). By contrast, a probe that is strongly cationic (e.g. 8K-Farn, 8+) is greatly stabilized in highly anionic membranes such as the PM and is not found on compartments of intermediate electronegativity. We therefore reasoned that if PS contributes to the electrostatic properties of intracellular compartments, it should accumulate in compartments that are electronegative. To test this idea, we crossed the mCITRINE^4K^4^Q-Farn^ (4+) reporter with the 2xmCHERRY-C2^LACT^ sensor and confirmed that both probes colocalized (Figure 5C). In addition, we found that 2xmCHERRY-C2^LACT^ colocalized preferentially with mCITRINE^4K^4^Q-Farn^ (4+) (which labels electrostatic compartments) rather than mCITRINE^8Q-Farn^ (0+) (which localization is charge independent) (Figure 5D). We next performed quantitative colocalization assay between intracellular compartment markers and charge reporters containing a gradual increase in net positive charges (0+, 2+ and 4+) in order to test the relative contribution of their positive charges on their intracellular distribution. We did not use probes with higher net positive charges than 4+, because the mCITRINE^6K^2^Q-Farn^ (6+) seldom localizes in intracellular compartments and mCITRINE^8K-Farn^ (8+) is strictly localized at the PM (Simon et al., 2016). We found that addition of positive charges gradually increased the proportion of the probes in EE/TGN and SV at the expense of their Golgi and late endosomes localization (Figure 5E). Therefore, the endomembrane system is organized according to an electrostatic gradient that is the highest at the PM, intermediate in early endocytic compartments, and low in the Golgi and late endosomes. This electrostatic gradient correlates with the PS concentration gradient, which suggests that PS might be involved in defining this electrostatic territory.

### Membrane surface charge probes relocalize to PS-bearing organelle in the absence of PI4P

Next, to grasp whether intracellular PS could control the electrostatic properties of intracellular membrane compartments, we inhibited PI4P synthesis using a 30 minutes PAO treatment (Simon et al., 2016) and asked where the mCITRINE^8K-Farn^ (8+) relocalized inside the cell. Interestingly, following PAO treatment, mCITRINE^8K-Farn^ was observed on the surface of PS bearing organelles, being mainly localized in early endocytic compartments and to a lower extent in late endosomes (Figure 6A-B and S6A-C). These results suggest that in the absence of PI4P, which is required for the distinctively high PM electrostatic signature (Simon et al., 2016), strongly cationic membrane surface charge reporters (such as the mCITRINE^8K-Farn^ reporter) localize inside the cell according to the PS concentration gradient.

**Figure 6.**
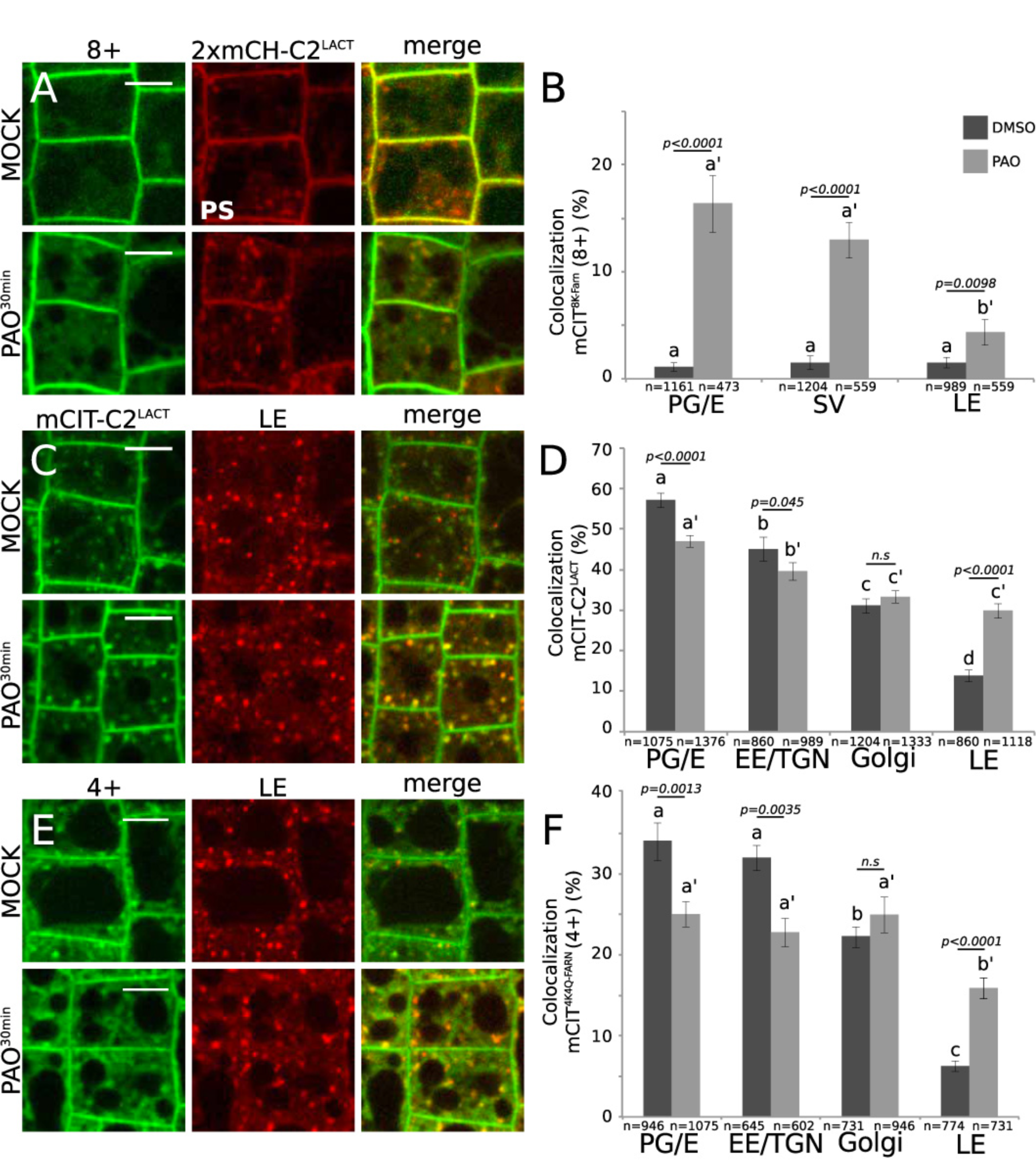
PS and PI4P cooperate to control endosome electrostatics. **A,** Confocal images of plants co-expressing mCITRINE^8K-Farn^ (8+) with mCHERRY-C2^LACT^ in mock (top) and PAO (60µM, 30 min, bottom) treated conditions. **B,** Quantification (mean ±s.e.m) of the percentage of compartments labelled by mCITRINE^8K-Farn^ (8+) that also contain W25R (PG/E), W24R (SV), and W7R (LE) in presence or absence of PAO (60µM, 30 min) n=(478, 1204) cells **C,** Confocal images of plants co-expressing mCITRINE-C2^LACT^ with W7R (LE) in mock (top) and PAO (60µM, 30 min, bottom) conditions. **D,** Quantification (mean ±s.e.m) of the percentage of compartments labelled by mCITRINE-C2^LACT^ that also contain W25R (PG/E), W13R (EE/TGN), W18R (Golgi), and W7R (LE) in presence or absence of PAO (60µM, 30 min), n=(860, 1376) cells. **E,** Confocal images of plants co-expressing mCITRINE^4K^4^Q-Farn^ (4+) with W7R (LE) in mock (top) and PAO (60µM, 30 min, bottom) conditions. **F,** Quantification (mean ±s.e.m) of the percentage of compartments labelled by mCITRINE^4K^4^Q-Farn^ (4+), that also contain W25R (PG/E), W13R (EE/TGN), W18R (Golgi), and W7R (LE) in presence or absence of PAO (60µM, 30 min) n=(602, 1075) cells. In graph B, D and F, different letters indicate significant differences among means (normal letters for DMSO comparison and letters with a prime symbol for PAO comparison, p value=0.05, Kruskal-Wallis bilateral test). Statistical difference between each sample is indicated by the p value at the top of each compared conditions (p-value=0.05, non-parametric Wilcoxon-Mann-Whitney test, non-significant (n.s.)). “n” represents the estimated number of cells sampled in each condition. Scale bars, 5 µm.

We previously noticed that PI4-kinase inhibition by PAO affects PS intracellular distribution (Simon et al., 2016). We therefore analyzed quantitatively PS subcellular localization in the absence or presence of PAO. We found that PAO treatment attenuated the gradient of PS as visualized by mCITRINE-C2^LACT^ (Figure 6D). In particular, PAO treatment increased the localization of mCITRINE-C2^LACT^ in late endosomes (Figure 6C-D). Strikingly, the electrostatic gradient, as visualized by the mCITRINE^4K^4^Q-Farn^ (4+) charge reporter was similarly affected by PAO treatment, with an increased localization of the reporter in late endosomes (Figure 6E-F and S6D-G). These results further confirm that charge reporter localization coincides with the presence of PS at the surface of intracellular membranes and support the notion that PS contributes overall to the establishment of the plant electrostatic territory at the surface of the PM cytosolic leaflet and along the endocytic pathway. In addition, we also noticed that PAO treatment decreased the accumulation of the mCITRINE^4K^4^Q-Farn^ (4+) probe in early endosomes (Figure 6F), while PS localization in this compartment was only mildly affected by this treatment (Figure 6D). PAO affects PI4P production, a lipid that is present in EE/TGN albeit to a lower extent than the PM (Simon et al., 2016). Loss of PI4P may therefore impact the electrostatic properties of EE and may explain the decreased accumulation of the mCITRINE^4K^4^Q-Farn^ (4+) probe in this compartment. As such, PI4P likely acts in combination with PS to specify the intermediate electronegativity of EE/TGN.

## Discussion

Here, we addressed which organelles are found in the electrostatic territory in plants and what are the anionic lipids that control this territory. Similar to previously published models, we found that the plant electrostatic territory corresponds to PM-derived organelles (Bigay and Antonny, 2012; Jackson et al., 2016). However, interestingly, we noticed that not all membranes in this territory are equally anionic. Rather, we revealed the existence of an electrostatic gradient, which is at its highest at the PM, intermediate in early endosomes and low in late endosomes. This electrostatic gradient is set up by various anionic phospholipid combinations. The concomitant accumulation of PA, PS and PI4P drives the very high electrostatic field of the PM. However, PS accumulation extends beyond the PM as it accumulates along the endocytic pathway according to a concentration gradient. This PS cellular distribution resembles that of animal cells, and contrast to that of yeast, in which PS massively accumulates at the PM (Moravcevic et al., 2010; Moser von Filseck et al., 2015a; Yeung et al., 2008). Furthermore, like in animals, the PS subcellular distribution in plants closely matches the electrostatic gradient, suggesting that PS is likely instrumental in setting up the electrostatic territory. In this scenario, PS and PI4P, which are found in the EE/TGN, drive the intermediate electrostatic property of this compartment. However, PS is also present in LE, where it may contribute to the weak electrostatic field of the late endocytic pathway. Phosphatidylinositol-3-phosphate (PI3P) and phosphatidylinositol 3,5-bisphosphate (PI(3,5)P_2_), are also enriched in LE (Noack and Jaillais, 2017), but are extremely rare lipids, which is consistent with the weak electronegativity of these compartments.

### PS as a general landmark of electrostatic membranes

The idea of two membrane territories, with distinct lipid compositions, as a fundamental organizing principle of the endomembrane system of eukaryotic cells was first proposed by Antonny and colleagues (Bigay and Antonny, 2012). These two lipid territories correspond roughly to ER and PM-derived membranes, and are defined by opposite physicochemical parameters (Bigay and Antonny, 2012; Jackson et al., 2016). The cytosolic leaflet of ER derived membranes is characterized by its low electrostatic property (as the vast majority of anionic phospholipids in the ER are orientated toward the lumen) and by its high occurrence of lipid packing defects, which are promoted by unsaturated lipids and the presence of small lipid head groups (Bigay and Antonny, 2012). By contrast, PM-derived organelles have few packing defects but are electrostatic, as they accumulate anionic phospholipids. PS is localized in PM-derived organelles in mammalian cells and may thereby contribute to the electrostatic properties of these compartments (Yeung et al., 2008). However, the importance of PS in mediating membrane surface charges along the animal endocytic pathway was deduced from pharmacological approaches that are known to also affect other cellular lipids (Ma et al., 2017; Yeung et al., 2008). Here, we combined pharmacological and genetic approaches to demonstrate that in plants PS is both necessary to establish the PM electrostatic signature and sufficient to maintain a certain degree of surface charges at the PM. Thus our results further consolidate the notion that PS is an important lipid across eukaryotes to establish the electrostatic territory (Jackson et al., 2016; Platre and Jaillais, 2017). However, by contrast to the proposed model, we further demonstrated that PS does not act alone in this process but rather do so in concert with PI4P and PA.

### Plants cells have significant PA levels in their plasma membrane, which is required to maintain the electrostatic properties of the PM cytosolic leaflet

It is well established that PA acts as a lipid messenger in plants, notably in response to the environment (Testerink and Munnik, 2011). In fact, almost every environmental stress triggers PA production within minutes, including abiotic stresses (e.g. cold, heat, drought, wounding, salinity) and biotic interactions (Testerink and Munnik, 2011). This rapid induction happens mostly at the PM and is regulated by direct production of PA by Phospholipase D (PLD) and/or by diacylglycerol phosphorylation by DGKs (Testerink and Munnik, 2011). Interestingly, in the present study, we found that the plant PM has significant PA level, as visualized by the recruitment of PA-binding sensors, even when plants are grown in optimal conditions. Plasma membrane recruitment of a PA reporter was previously observed in sub-domain of tobacco pollen tubes plasmalemma (Potocky et al., 2014) and seems to be extendable to most of the tissues we observed in *Arabidopsis*. In animals, most cells have minute amount of PA at the PM and PA sensors are not recruited to the PM in resting conditions (Bohdanowicz et al., 2013). By contrast, phagocytic cells, such as macrophages and immature dendritic cells, have relatively high level of PA in their PM (Bohdanowicz et al., 2013). This unusual concentration of PA allows these cells to have constitutive membrane ruffling in order to scan their environment, which is required for immune surveillance. These phagocytic cells maintain their elevated PA level at the PM via DGK activity (Bohdanowicz et al., 2013). Similarly, we found that in plants a DGK activity is required to sustain the level of PA at the PM. Pharmacological inhibition of DGK activities not only solubilizes PA sensors but also impacts PM electrostatic properties. Using similar approaches, it was recently shown that PA plays a role in the PM targeting of the D6-PROTEIN KINASE (D6PK) (Barbosa et al., 2016), an AGC kinase involved in polar auxin transport (Armengot et al., 2016). The localization of D6PK is dependent on both PI4P, PI(4,5)P_2_ and PA, suggesting that a combination of phosphoinositides and PA is responsible for its localization rather than a single phospholipid species (Barbosa et al., 2016). Here, we obtained similar results with several independent generic membrane surface charge reporters, suggesting that the requirement for several anionic phospholipids may not be an intrinsic property of D6PK but rather a more general feature of the electrostatic field of the plant PM. In addition, this further suggests that our results are not just limited to our synthetic charge reporters, but are relevant for the localization of endogenous *Arabidopsis* proteins, and point toward a more general requirement of PI4P/PA/PS combination for the localization of many proteins in plants. However, it is worth mentioning that the localization of D6PK is not identical to that of generic membrane surface charge sensors. Indeed, unlike these sensors, D6PK localizes at the basal pole of the cell, and its PM targeting is highly BFA sensitive (Barbosa et al., 2014; Simon et al., 2016). It is therefore likely that D6PK localization is regulated by additional factors, which might include PI(4,5)P_2_ as well as other, yet unknown regulators. Mutations of the cationic residues responsible for anionic lipid interactions in the middle region of D6PK do not completely abrogate its membrane binding, suggesting that indeed D6PK localization is not solely regulated by electrostatic interactions with anionic phospholipids (Barbosa et al., 2016). Consistently, D6PK N- and C-terminal end are also required for D6PK localization (Barbosa et al., 2016).

In yeast and animal cells, PM electrostatics is extremely robust, with the loss of one anionic phospholipid species having little or no impact on the overall charge of the PM. By contrast in plants, we found that the individual loss of PI4P, PS and PA directly impact PM electrostatics. While they are all anionic phospholipids, they have radically different turnover. Indeed, PS is a relatively stable phospholipid, while PI4P and PA have a high turnover rate. One may speculates that PS ensures a stable PM electrostatic field, while spatiotemporal variations in PI4P and/or PA may directly impact PM surface charges. As such, PM electrostatics in plants may be particularly prone to respond to environmental changes. It will be an exciting future direction to understand how environmental stresses impact membrane electrostatics, what are the contributions of individual lipids in these variations and how this might impact signaling, intracellular trafficking and cellular polarity.

## Acknowledgements

We thank A. Martinière-Delaunay, O. Hamant, M. Dreux and the SiCE group for discussions and comments, G. Du, T. Taguchi, S. Grinstein, B. Kost and addgene for providing plasmids, N. Geldner, K. Schumacher, I. Moore, D. Ehrhardt, the GABI-KAT, and the NASC collection for providing transgenic *Arabidopsis* lines, A. Lacroix, J. Berger and P. Bolland for plant care, J.C. Mulatier for help in preparing lipids and PLATIM for help with imaging. Y.J. is funded by ERC no. 3363360-APPL under FP/2007-2013. M.D. and M.S are funded by a PhD fellowship from the French Ministry of Higher Education. T.S is supported by ERC grant no. 615739-MechanoDevo to O. Hamant. P.P and M.P. are supported by the Czech Science Foundation grant no. 17-27477S. Lipidomic analyses were performed on the Bordeaux Metabolome Facility-MetaboHUB (ANR-11-INBS-0010).

## Author contributions

M.P.P. was responsible of all experiments described in the manuscript except the following: production and imaging of the *PDF1*-driven PS reporter line (T.S.), lipid measurements (L.M-P., L.F. and P.M.); time-lapse imaging of cytokinesis/root hair (M.D. and M.C.C.) and pollen tube (P.P. and M.P.). V.B. helped with image quantification and acquisition, L.A. helped with yeast and lipid overlay experiments; A.C. helped with LPS/LPA addback experiments, M.S. participated in the production of membrane charge reporter lines. M.P.P., and Y.J. conceived the study, designed experiments and wrote the manuscript and all the authors discussed the results and commented on the manuscript.

---

**Figure S1.**
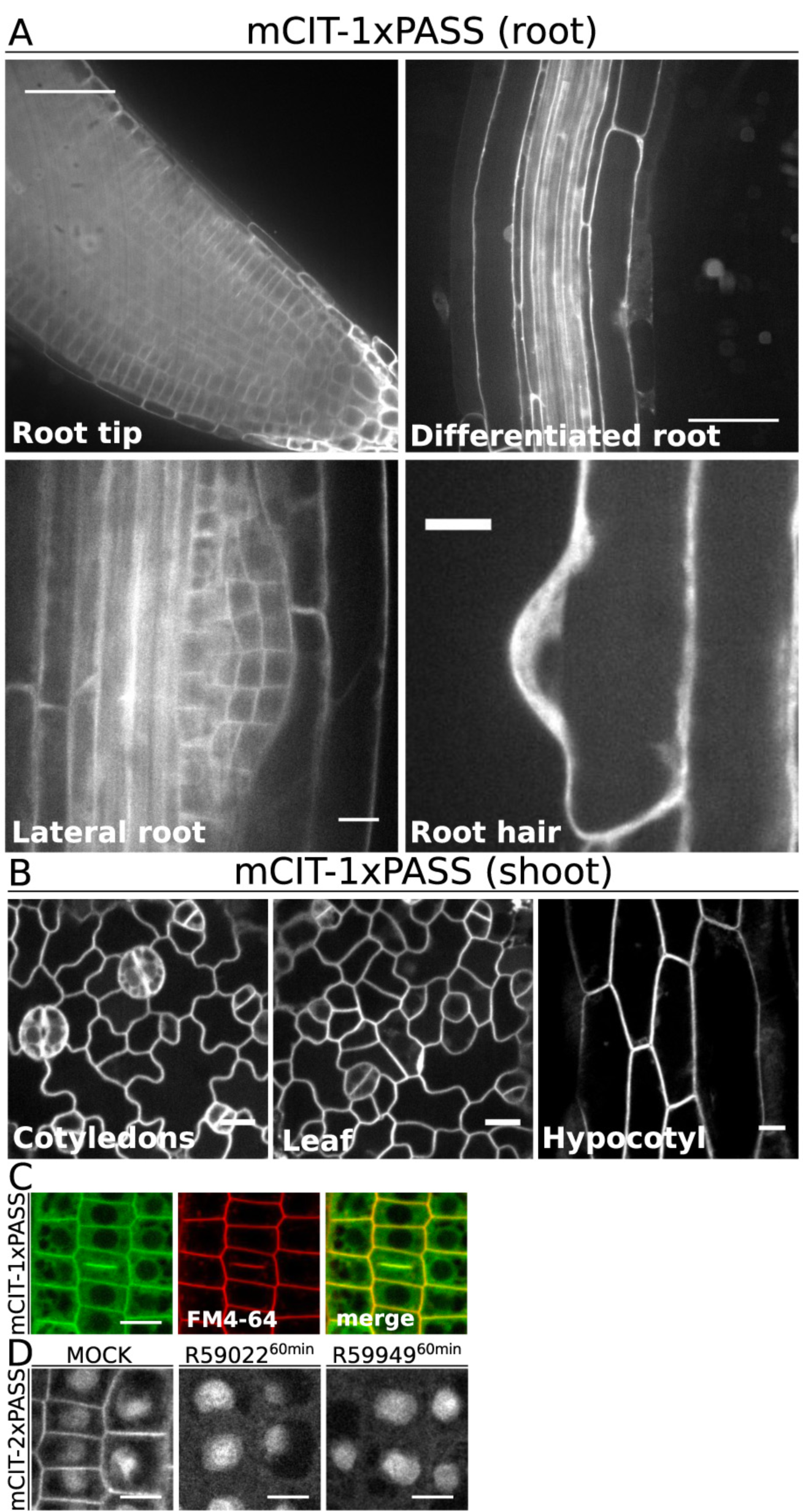
(Related to Figure 1). PA sensors localize at the plasma membrane in different cell types. **A**, Confocal images of plants expressing mCITRINE-1xPASS in different root tissues: root tip (top left), differentiated cells (top right), lateral root primordium (bottom left) and bulging root hair (bottom right). **B**, Confocal images of plants expressing mCITRINE-1xPASS in different shoot tissues: cotyledons (left), leaf (middle) and hypocotyl (right). **C**, Confocal images of *Arabidopsis* root epidermis stained by FM4-64 (1µM, 60min) and expressing mCITRINE-1xPASS showing co-labelling at the cell plate. **D**, Confocal images of plant expressing mCITRINE-2xPASS in control condition (right), or following DGK inhibition by R59022 (12.5µM, 60 min, middle) or R59949(12.5µM, 60 min, right).

**Figure S2.**
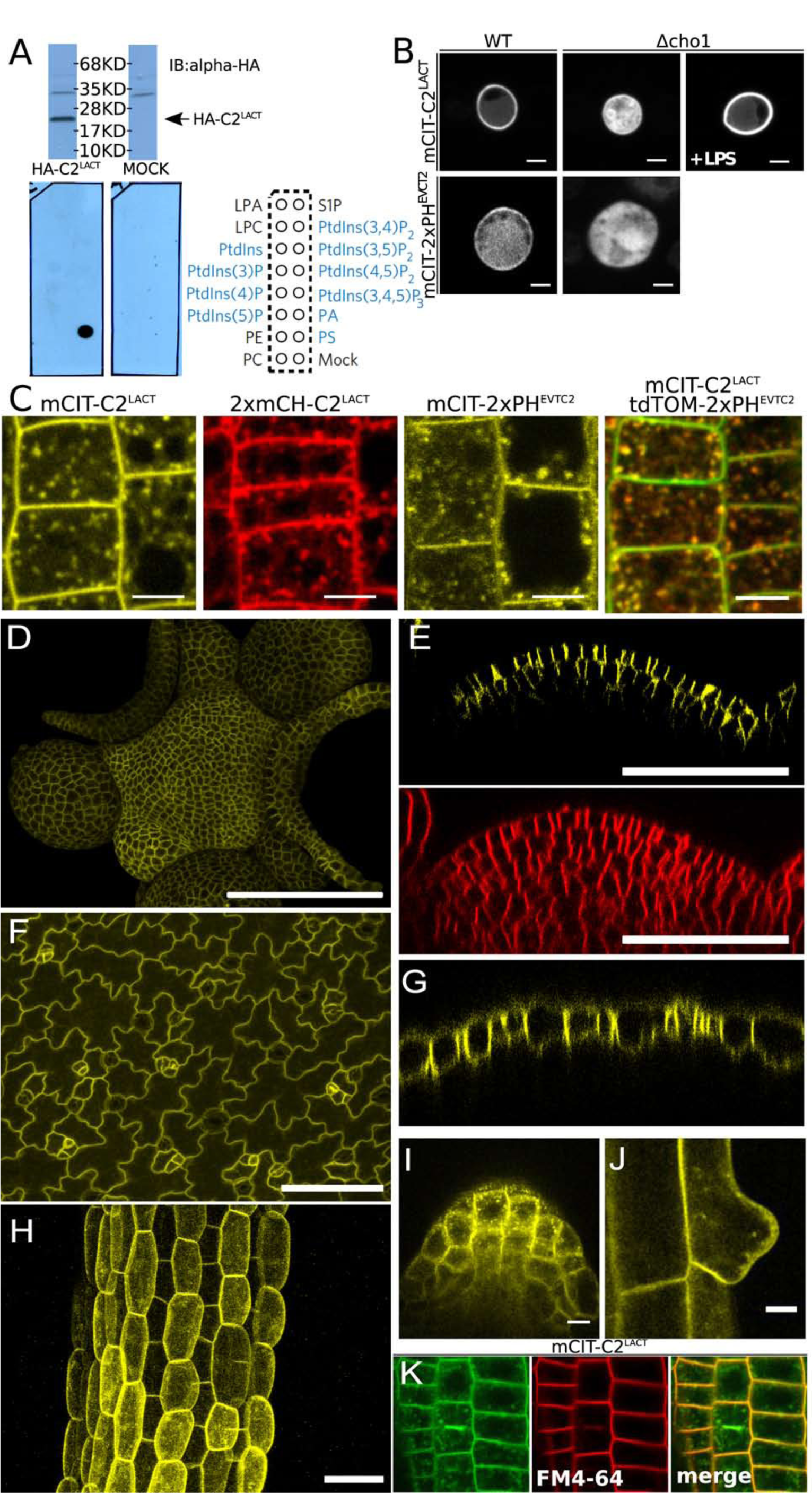
(Related to Figure 2). Characterization of PS sensors localization in different cell types. Although the C2 domain of Lacthaderin has been extensively used as a PS reporter, we verified the PS-binding selectivity of our construct, which differs from published reporters in its linker sequence between the C2^LACT^ domain and the fluorescent proteins. All our constructs were obtained using recombination-based cloning. We can therefore switch tags and expression systems while keeping the linker sequence constant. First, we found that in vitro translated HA-C2^LACT^ specifically binds to PS in lipid-protein overlay assays (Fig. S2A), confirming previous binding assays performed with liposomes (Yeung et al., 2008). Second, we tested the localization and PS sensitivity of our C2^LACT^ construct in vivo using recombinant expression in wild type and *cho1Δ* yeast strains, the latter being deficient for PS biosynthesis (Fig. S2B). As we previously reported (Simon et al., 2016), our C2^Lact^-GFP construct localizes at the plasma membrane (PM) in WT yeasts and is soluble in the absence of PS in the *cho1Δ* mutant (Fig. S2B). In addition, the soluble localization of C2^LACT^ in *cho1Δ* is rescued by one hour of exogenous treatment with lysoPS (LPS), confirming that our construct behaves as previously described C2^LACT^ probes (Maeda et al., 2013; Moser von Filseck et al., 2015b; Yeung et al., 2008). Together, these results validate the PS-selectivity of our C2^Lact^ construct. Furthermore, we verified that it colocalizes with another PS binding protein, the PH domain of human EVECTIN2 (PH^EVCT2^), which has also been used as a PS reporter in vivo. In *Arabidopsis* root epidermis, mCITRINE-2xPH^EVCT2^ showed a similar localization pattern as the C2^LACT^ reporter and mCHERRY-2xPH^EVCT2^ extensively colocalizes with mCITRINE-C2^LACT^ (Fig S2C). Similar to C2^LACT^, we validated our PS PH^EVCT2^ probe specificity using heterologous expression in WT and *cho1Δ* mutant yeast (Fig. S2B). Together, these approaches validated C2^LACT^ as a *bona fide* PS reporter in plants. **A,** Western blot showing expression of recombinant HA-C2^LACT^ (top), lipid overlay assay performed with HA-C2^LACT^ (bottom left), empty vector (bottom middle) and scheme showing the position of the different lipid species spotted on the membrane (bottom right), anionic lipids are highlighted in blue. **B**, Confocal images of yeast expressing GFP-C2^LACT^ upper panel and GFP-1xPH^EVCT2^. Left pictures correspond to wild type background, middle to *Δcho1* yeast strain depleted of PS and right *Δcho1* yeast strain complemented with LPS (54µM 60 min). Scale bars, 5 µm. **C**, Confocal images of plants expressing PS sensors. From left to right, mCITRINE-C2^LACT^, 2xmCHERRY-C2^LACT^ and mCITRINE-2xPH^EVCT2^ and plants co-expressing, 2xmCHERRY-2xPH^EVCT2^ with mCITRINE-C2^LACT^. Scale bars, 5 µm. **D-H**, Plant expressing mCITRINE-C2^LACT^ driven by the shoot- and L1-specific *PDF1* promoter in different shoot tissues. **D**, top view of the shoot apical meristem, **E**, Cross-section in the central zone of the shoot apical meristem (top) and FM4-64 staining for 60 min (bottom), **F**, cotyledon epidermis and **G**, a cross-section in cotyledons epidermis **H**, Z-projection of z-stacks taken in the hypocotyl. **I-J**, Confocal images of UBQ10prom::mCITRINE-C2^LACT^ in lateral root primordium (**I**) and in bulging root hair (**J**). **K**, Confocal images of *Arabidopsis* root epidermis stained by FM4-64 (1µM, 60min) and expressing mCITRINE-C2^LACT^ showing co-labelling at the cell plate. Scale bars, 5 µm.

**Figure S3.**
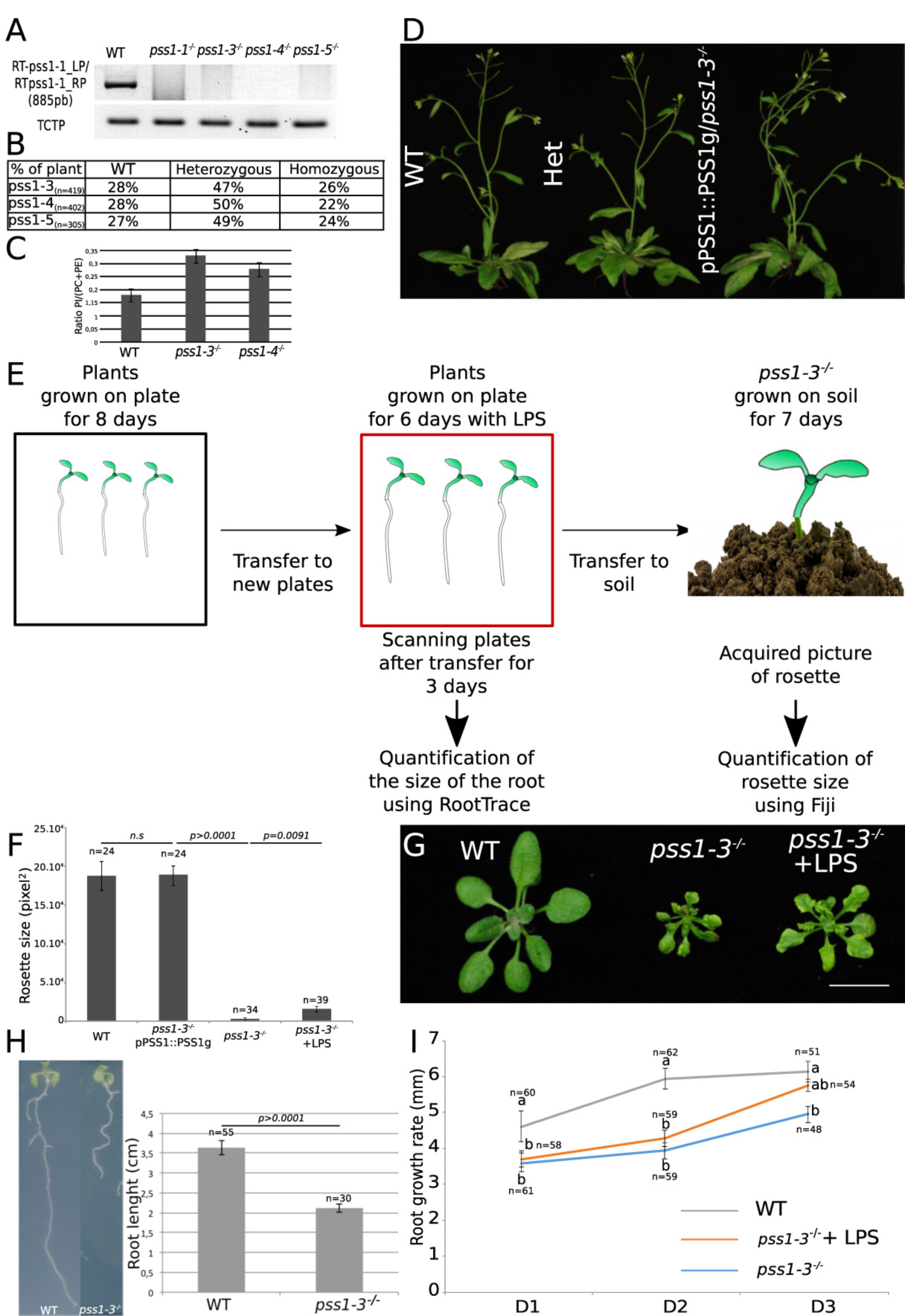
(Related to Figure 3). Characterization of *pss1* mutants. **A,** RT-PCR analysis of *PSS1* transcript in WT and *pss1* mutant showing the absence of full length *PSS1* transcript in *pss1-1* to *pss1-4* alleles. The bottom row shows expression of ubiquitously express *TCTP* gene in both WT and *pss1* mutants. **B**, Segregation analysis from *pss1* heterozygous plants for *pss1-3, pss1-4* and *pss1-5*, in percentage. **C,** Quantification of the ratio of PI/(PC+PE) in WT and *pss1* mutants. This ratio was obtained by measuring the area bellow the pics corresponding to PI, PE and PC for each genotype (WT, n=6; *pss1-3*, n=8 and *pss1-4*, n=8). This analysis shows that *pss1* mutants have a slight elevation in their total PI content at the expense of PC and PE. **D**, Comparison of 45 day-old plants between a wild type plant (left), a *pss1-3*^*+/-*^ heterozygous plant (Het, middle) and *pss1-3*^*-/-*^ homozygous plant complemented by transgenic expression of a *PSS1* genomic fragment (*pPSS1::PSS1g*). **E,** Schematic representation of the procedure to complement plants with LPS in order to quantify the root growth rate and the rosette area. **F,** Quantification of the rosette area (mean ±s.e.m in pixel^2^) of wild type plants, *pss1-3*^*-/-*^ mutants expressing *pPSS1::PSS1g, pss1-3*^*-/-*^ mutants and *pss1-3*^*-/-*^ mutants treated with exogenous LPS at 2.47µM. Statistical difference between each sample is indicated by the p value at the top of each compared conditions (p-value=0.05, non-parametric Wilcoxon-Mann-Whitney test, non-significant (n.s.)). “n” correspond to the number of plants used. **G**, Picture showing the rosette of 21-day-old wild type plants, *pss1-3*^*-/-*^ and *pss1-3*^*-/-*^ supplemented with LPS for 6 days. (see Fig S3G). **H**, Picture showing 12 days-old seedlings of wild type (left) and *pss1-3*^*-/-*^ (right) plants. Statistical difference between each sample is indicated by the p value at the top of each compared conditions (p-value=0.05, non-parametric Wilcoxon-Mann-Whitney test). “n” correspond to the number of plants used. **I**, Quantification (mean ±s.e.m in mm) of root growth for 3 days in wild type, *pss1-3*^*-/-*^ and *pss1-3*^*-/-*^ supplemented with LPS at 2.47µM. D1-D2-D3 correspond to one, two or three days after LPS treatment, respectively. Different letters indicates statistical difference between samples (p value=0.05, Kruskal-Wallis bilateral test). “n” correspond to the number of plants used.

**Figure S4.**
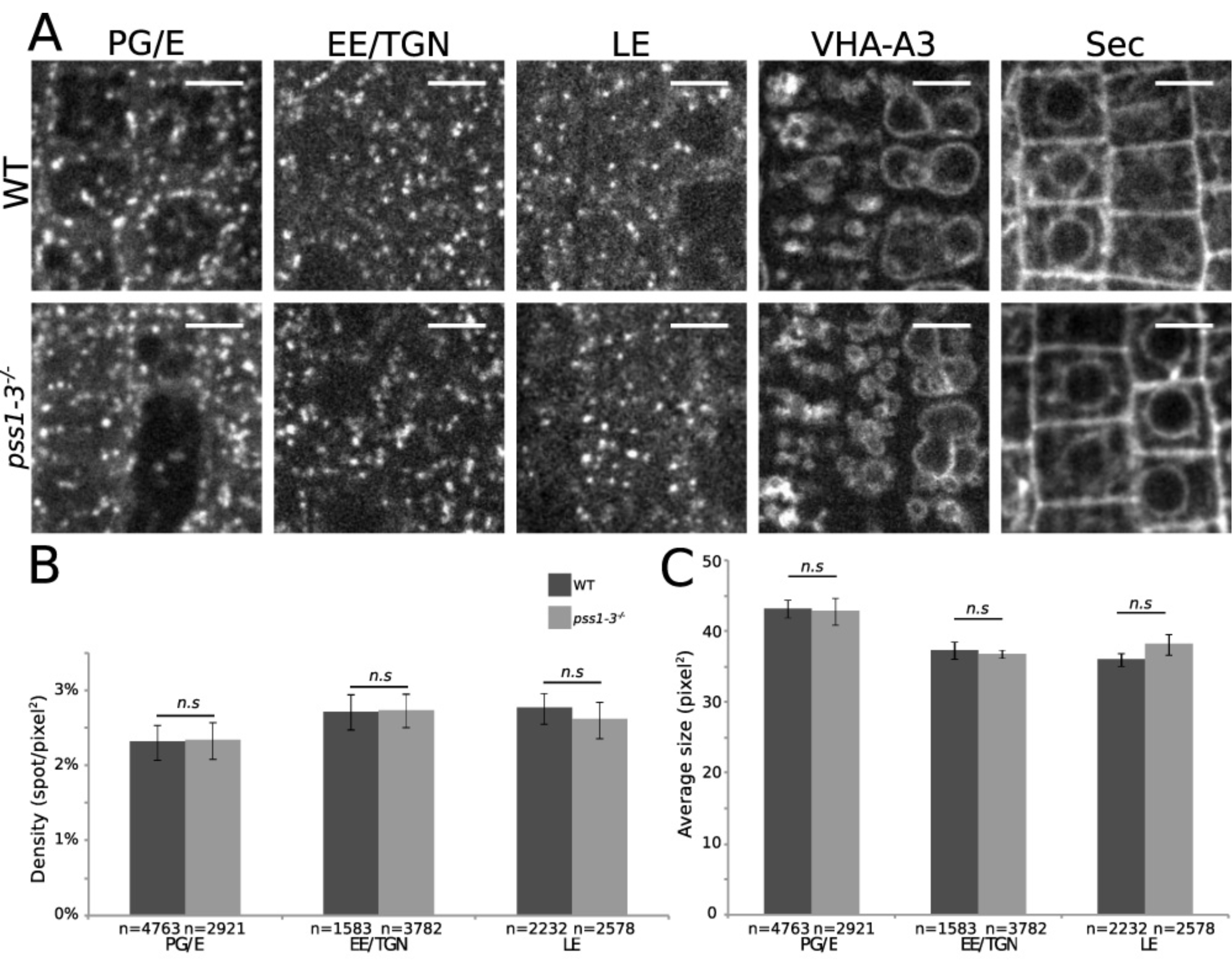
(Related to Figure 4). Intracellular compartmentalization is not affected in *pss1-3*^*-/-*^. **A,** From the left to the right, plants expressing W25R (post-Golgi/endosomal (PG/E)), W13R (Early endosomes/*trans*-Golgi network (EE/TGN)), W7R (Late endosomes (LE)), VHA-A3-RFP (tonoplast), and Sec-RFP (secretion) in wild type plant (upper panel) and in *pss1-3*^*-/-*^ (lower panel). **B,** Quantification (mean ±s.e.m, number of spots per pixel^2^) of the density of intracellular compartments labeled by W25R (PG/E), W13R (EE/TGN), W7R (LE) in wild type and *pss1-3*^*-/-*^**. C,** Quantification (mean ±s.e.m, size in pixel^2^) of the average size of intracellular compartments labeled by W25R (PG/E), W13R (EE/TGN), W7R (LE) in wild type and *pss1-3*^*-/-*^. Statistical difference between each sample is indicated by the p value at the top of each compared conditions (p-value=0.05, non-parametric Wilcoxon-Mann-Whitney test, non-significant (n.s.)). “n” represents the number of spots sampled in each condition. Scale bars, 5 µm

**Figure S5.**
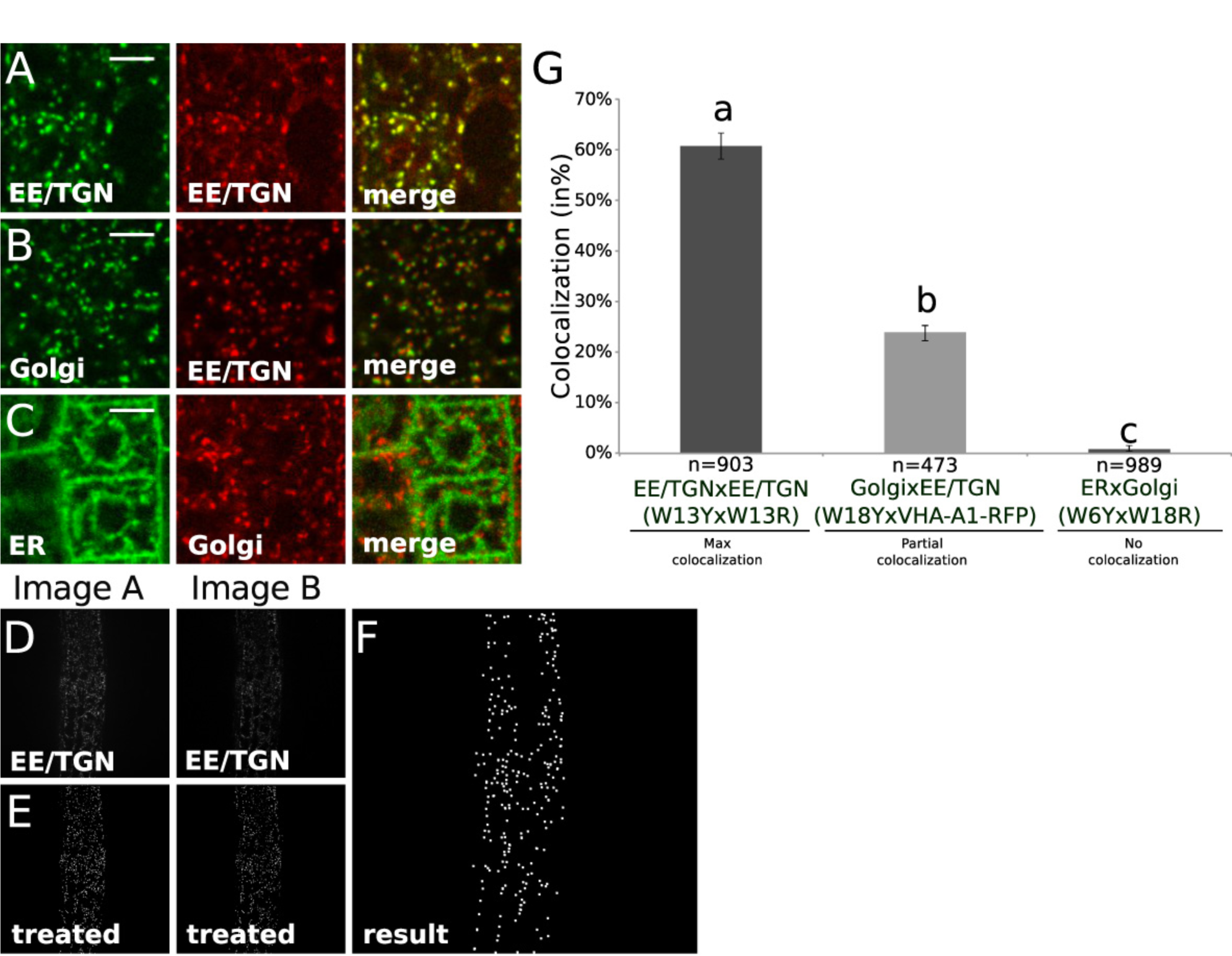
(Related to Figure 5). Validation of the quantitative colocalization methods used in this study. **A,** Confocal images of plant co-expressing EE/TGN marker W13Y (left) and W13R (middle) and the corresponding merge (right). **B**, Confocal images of plant co-expressing Golgi marker W18Y (left) and early endosomal marker VHA-A1-RFP (middle) and the corresponding merge (right). **C**, Confocal images of plant co-expressing endoplasmic reticulum marker W6Y (left) and Golgi marker W18R (middle) and the corresponding merge (right). **D**, Raw images of plant co-expressing EE/TGN marker W13Y (left, Image A) and W13R (right, Image B). **E**, Image processing applying a DoG filter with a sigma of 3 and a triangle thresholding for the corresponding image A (left) and B (middle). **F**, Each white spots indicate colocalization between spots issue from the treated image A and B. **G**, Quantification (mean ±s.e.m) of the percentage of colocalization of the indicated yellow wave line (WnY) with red wave line (WnR). Statistical analysis was performed using the non-parametrical Kruskal-Wallis test (p-value=0.05) and pairwise comparisons between groups was performed according to Steel-Dwass-Critchlow-Fligner procedure (a, b, c indicate statistical difference between samples). “n” represents the estimated number of cells sampled in each condition. Scale bars, 5 µm.

**Figure S6.**
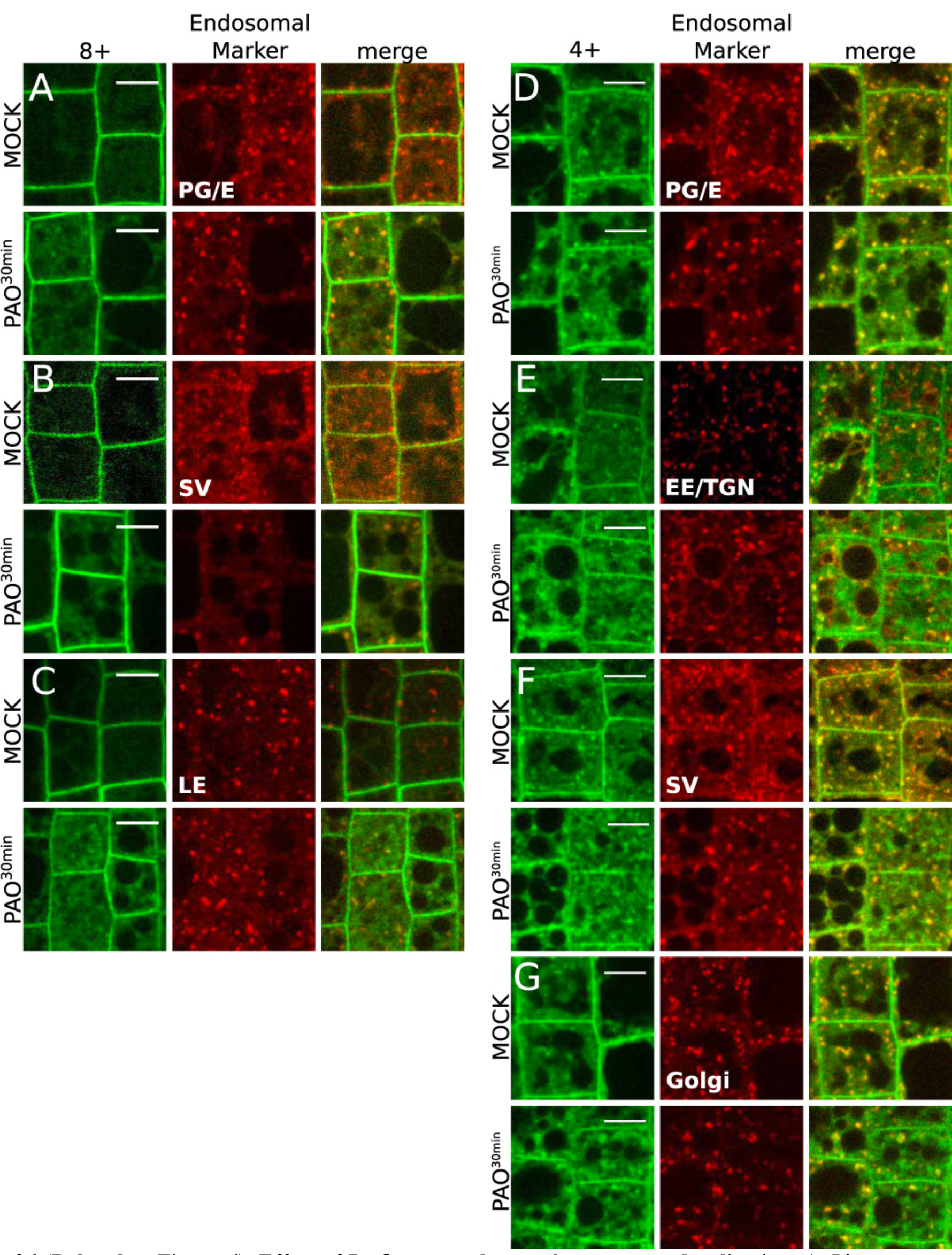
(Related to Figure 6). Effect of PAO on membrane charge sensor localization. **A**, Plant co-expressing mCITRINE^8K-Farn^ (left) and W25R post-Golgi/endosomal marker (middle), and the corresponding merge (right) in mock condition (top) and upon 60µM PAO treatment for 30 min (bottom). **B**, Plant co-expressing mCITRINE^8K-Farn^ (8+, left) and W24R secretory vesicle marker (middle), and the corresponding merge (right) in mock condition (top) upon 60µM PAO treatment for 30 min (bottom). **C**, Plant co-expressing mCITRINE^8K-Farn^ (8+, left) and W7R late endosomal marker (middle), and the corresponding merge (right) in mock condition (top) upon 60µM PAO treatment for 30 min (bottom).**D**, Plant co-expressing mCITRINE^4K^4^Q-Farn^ (4+, left) and W25R post-Golgi/endosomal marker (middle), and the corresponding merge (right) in mock condition (top) and upon 60µM PAO treatment for 30 min (bottom). **E**, Plant co-expressing mCITRINE^4K^4^Q-Farn^ (left) and W24R secretory vesicle marker (middle), and the corresponding merge (right) in mock condition (top) and upon 60µM PAO treatment for 30 min (bottom). **F**, Plant co-expressing mCITRINE^4K^4^Q-Farn^ (4+, left) and VHA-A1-RFP early endosomal marker (middle), and the corresponding merge (right) in mock condition (top) and upon 60µM PAO treatment for 30 min (bottom). **G**, Plant co-expressing mCITRINE^4K^4^Q-Farn^ (4+, left) and W18R Golgi marker (middle), and the corresponding merge (right) in mock condition (top panel) and upon 60µM PAO treatment for 30 min (bottom). Scale bars, 5 µm.

**Video S1. mCITRINE-1xPASS localizes on early cell plates**. Time lapse imaging of plant of 5 days old seedlings expressing mCITRINE-1xPASS in root epidermis. Images every 4 minutes. Scale bar, 5µm.

**Video S2. mCITRINE-2xPASS PA biosensor localizes at the plasma membrane in the flank region of root hair**. Time lapse imaging of plant of 5 days old seedlings expressing mCITRINE-2xPASS in the root hair. Images every 5 minutes. Scale bar, 5µm.

**Video S3. mCITRINE-C2^LACT^ PS biosensor localizes in the inverted cone of the root hair**. Time lapse imaging of plant of 5 days old seedlings expressing mCITRINE-C2^LACT^ in the root hair. Images every 5 minutes. Scale bar, 5µm.

**Video S4. PS biosensor localized in the inverted cone and to the subapical plasma membrane of the pollen tube**. Time lapse imaging of tobacco pollen tube transiently expressing YFP-C2^LACT^. Images were taken every 1 s. Movie is representative for 20 pollen tubes recorded in two independent experiments. Scale bar, 10 µm.

**Video S5. PI4P and PS biosensors are recruited concomitantly at the cell plate**. Time lapse imaging of plant of 5 days old seedlings co-expressing mCITRINE-1xPH^FAPP1^ and 2xmCHERRY-C2^LACT^ in root epidermis. Images every 3 minutes. Scale bar, 5µm.

